# AI methods and biologically informed data curation enable accurate RNA m^5^C prediction

**DOI:** 10.1101/2025.09.22.677824

**Authors:** Emanuele Saitto, Elena Casiraghi, Alberto Paccanaro, Giorgio Valentini

## Abstract

5-methylcytosine (m^5^C) RNA modifications influence nearly every aspect of RNA metabolism, but their transcriptome wide detection is limited by costly, error-prone assays. To bridge this experimental gap, a wave of AI tools now predicts putative m^5^C sites *in silico*. However, most existing approaches prioritize architectural complexity while neglecting data quality, so their reported gains mainly reflect the artifacts inherited from noisy datasets. We inverted this paradigm by constructing a high-confidence, methyltransferase-specific catalog of m^5^C sites, removing artifacts that confound existing resources. Using this curated corpus, we trained (for the first time in a multiclass setting) three different models (Bi-GRU, CNN, Transformer) to distinguish writer-specific m^5^C sites from unmethylated cytosines. All AI models converged to similar, nearly optimal, performance (AUPRC > 0.97), and a biologically informed analysis revealed that most errors clustered in unmethylated sites mimicking true positives. By augmenting the training set with these hard-to-predict negatives, mined from millions of unmodified cytosines, the models were forced to exploit more nuanced features such as RNA secondary structure and subtle sequence cues, which sharply reduced transcriptome-wide false positive predictions, and predicted methylated transcripts exhibited strong concordance with known methyltransferase biology. Explainable AI techniques also showed that our AI models effectively capture how sequence mutations disrupt m^5^C sites, underscoring their potential to prioritize disease-relevant variants. The main findings of our study underscore that AI models can be decisive levers for reliable m^5^C identification only if fed with curated data and validated through biologically informed computational analysis.

## 1 Introduction

RNA is not merely a linear polymer of four canonical nucleotides; more than 170 distinct covalent marks have been cataloged to date, expanding the chemical alphabet of the transcriptome well beyond A, C, G, and U.^1^ Together, these marks constitute the RNA epitranscriptome. By changing electrostatics, hydrophobicity and base-pairing, single-nucleotide modifications let cells fine-tune gene expression at every post-transcriptional step.^2–4^

Among the many RNA modifications, 5-methylcytosine (m^5^C) is a conserved post-transcriptional mark that influences virtually every aspect of RNA metabolism, including nuclear export, splicing, stability, and translation. Its effects are mediated both by changes in the physicochemical properties of RNA and by the recruitment of specific reader proteins (ALYREF, SRSF2, YBX1).^5–7^ m^5^C is installed by cytosine-5 RNA methyltransferases (mainly NSUN enzymes), which recognize short sequence motifs and local secondary structures around the target cytosine, thereby promoting writer binding and catalysis.^8^ Over evolutionary time, these sequence–structure contexts appear to have been selected to accommodate writer recognition, positioning m^5^C at sites where methylation can modulate RNA processing and function.

Notably, m^5^C has also been detected in viral genomes, highlighting its functional relevance beyond endogenous RNA.^9^ Conversely, dysregulation of m^5^C has been associated with neurological disorders, metabolic diseases, and various cancers.^8, 10^

Given its broad regulatory role and clinical relevance, uncovering the full extent of the m^5^C epitranscriptome is essential. To this end, several experimental techniques have been developed to detect RNA m^5^C modifications, including bisulfite sequencing (BS-seq), m^5^C-RNA immunoprecipitation (m^5^C-RIP), methylation-dependent individual-nucleotide resolution crosslinking and immunoprecipitation (miCLIP), and 5-azacytidine–mediated RNA immunoprecipitation (Aza-IP).^11–14^ Among them, RNA BS-seq is the most widely used technique for transcriptome-wide, base-resolution detection of m^5^C.^5, 11, 15–17^

Despite its success, BS-seq still faces significant challenges in achieving sensitive and accurate m^5^C detection.^15, 18, 19^ This shortcoming, as well as those of alternative laboratory assays (discussed in Suppl. Note 1) significantly obstruct our comprehension of m^5^C methylations, particularly in mRNAs, where the exact number of m^5^C sites continues to be a subject of debate.^5, 15, 19^

To overcome these limitations, machine and deep learning-based methods have been developed to predict transcriptome-wide m^5^C sites that remain beyond the reach of current laboratory techniques.^20–25^

While recent models represent a step forward in m^5^C site prediction, it is well-known that the performance of machine and deep learning models is highly dependent on data quality and selection.^26^ In the context of m^5^C prediction, most efforts have chased increasingly complex architectures^23–25^ while neglecting the quality of the training data. Because these datasets are derived directly from noisy, error-prone wet-lab assays, they contain substantial false positives, and models trained on them inevitably inherit such biases. Moreover, benchmarking on the same noisy datasets may lead to misleading performance estimates: an issue that has so far been masked by the absence of biological validation and model interpretation.

To overcome these limitations, we developed a computational framework that couples state-of-the-art deep learning with rigorous data curation and biologically informed validation, ensuring that model performance reflects true biological signal rather than artifacts of noisy datasets. Our main contributions can be summarized as follows:

- **Construction of a high-confidence training corpus:** Building on the extensive BS-seq catalog of Liu et al.,^27^ we developed a computational pipeline to construct a curated, writer-resolved m^5^C dataset. This involved methyltransferase-specific reclustering, false positive filtering and cluster refinement to mitigate BS-seq artifacts. Negative sites (unmodified cytosines) were sampled from the same mature transcripts containing at least one modification. This strategy avoided trivial discriminations (e.g., untranscribed DNA, intronic regions) while maximizing confidence in the negatives’ unmethylated status.
- **Design of accurate AI-based m**^5^**C predictors:** models based on Bidirectional Gated Recurrent Units (Bi-GRU), one-dimensional Convolutional Neural Networks (1D-CNN) and encoder-based Transformers trained on our curated dataset in a multiclass setting achieved accurate predictions, and were able to confidently distinguish with high accuracy (AUPRC > 0.97) the methyltransferase classes underlying the m^5^C modifications.
- **Model refinement, interpretation and biologically informed validation:**
  – *Hard-negative mining*: our AI models retrained with enriched “hard negatives” (i.e. the most difficult to predict unmodified cytosines), not only better predicted transcription-wide m^5^C sites, but were also able to restore experimentally supported nuanced features such as writer-specific RNA secondary structure and writer-specific motifs overlooked by other models.
  – *Model interpretation and variant effect modeling:* An interpretable surrogate model explains m^5^C predictions of our AI models, and reveals the mutational effects on m^5^C sites learned by them, underscoring their potential to prioritize disease-relevant variants.
  – *Functional validation*: Gene enrichment analysis revealed novel predicted m^5^C sites mapping to genes with functions consistent with known methyltransferase-specific biology.

We release all code, model parameters, training and evaluation scripts, and datasets. In addition, we provide a lightweight prediction tool (FASTA input; csv/tsv/xlsx output) and a transcriptome-wide, writer-specific catalog of high-confidence predicted m^5^C sites for community use.

## 2 Results

The first step in our work was to select available m^5^C data for training predictors. A survey of the literature revealed that some resources report only a limited number of methylation calls, and more generally, many lack rigorous validation. Considering the high error rate of wet-lab assays (mentioned above and discussed in detail in Suppl. Note 1), reported m^5^C sites without proper validation remain of questionable reliability. A major breakthrough in this context came with the work of Liu et al.,^27^ who applied an optimized RNA BS-seq protocol to profile oocytes and early embryos across six animal phyla. This approach uncovered unexpected *waves* of maternal m^5^C, a developmental stage absent in adult tissues. The breadth of the dataset enabled more stringent filtering of false positives while preserving a large number of confident calls.

Given that this was the most extensive and validated resource, we chose it as the foundation for training our models. The overall framework of our work is summarized in Fig. 1. First, we constructed a high-confidence catalog of human m^5^C sites by rigorously filtering the Liu et al.^27^ dataset to remove residual bisulfite artifacts and clustering sites to enforce methyltransferase specificity (Fig. 1a). Unbiased negatives were retrieved from the same mature transcripts that harbored at least one methylation call, providing the strongest confidence in their unmethylated status.

**Figure 1:**
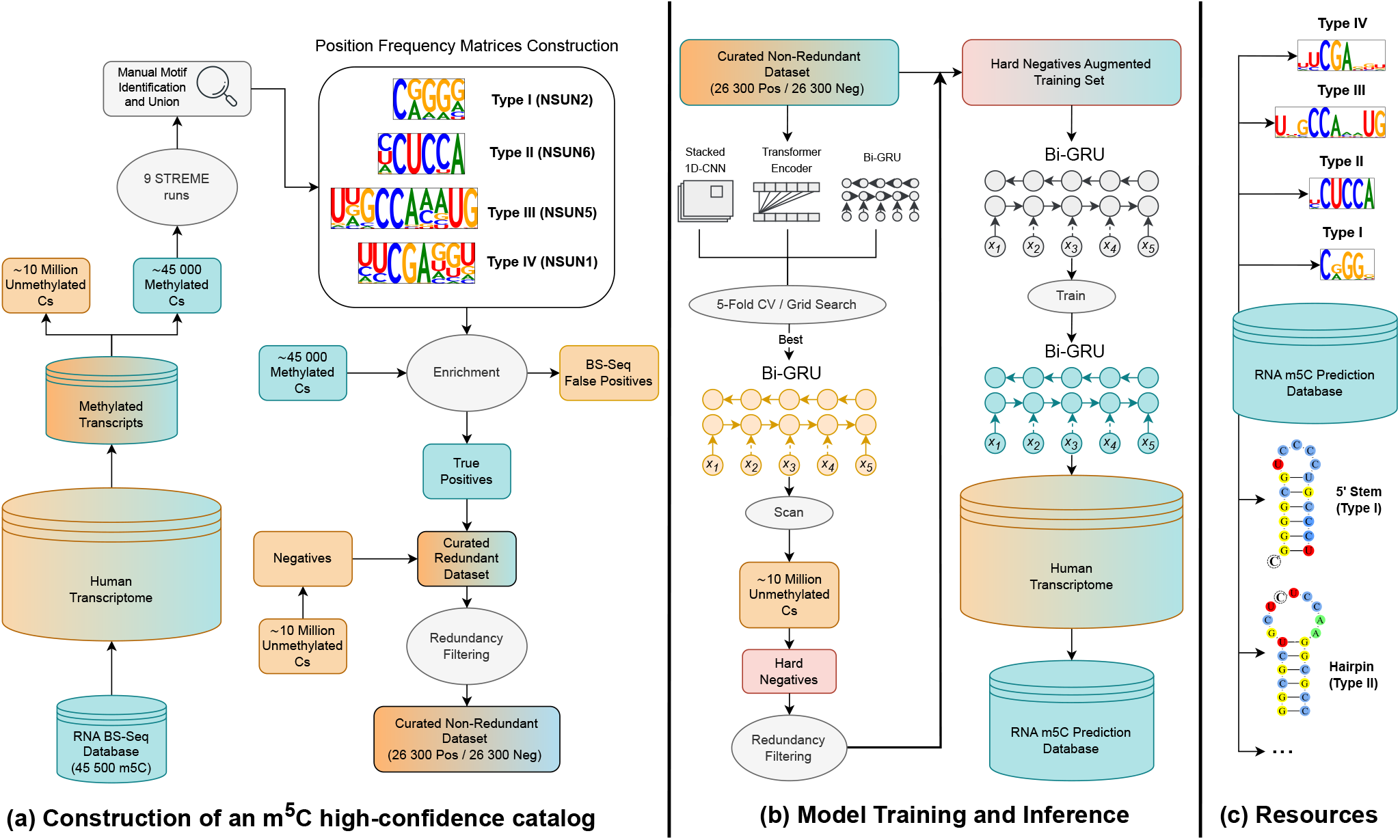
End-to-end pipeline for writer-resolved m^5^C prediction. **(a) Construction of a high-confidence m**^**5**^**C catalog**. The 45 500 bisulfite calls of Liu et al.^27^ are mapped to GENCODE v19 transcripts, and cytosines on methylated mature transcripts are partitioned into methylated (turquoise) and unmethylated (orange). To recover methyltransferase-specific sequence patterns, a motif-discovery step (STREME) is applied to the short sequence context surrounding each methylated cytosine; because this procedure is stochastic, it is repeated across multiple independent runs and recurring patterns are manually merged to obtain four robust writer motifs (Types I–IV, boxed). Motif-derived position-frequency matrices are then used to rescore all candidates, removing residual BS-seq artifacts and assigning sites to four writer-specific clusters. Adding transcript-matched negatives and applying redundancy filtering yields a non-redundant 26 300×2 corpus. (b) **Model Training and Inference**. A five-fold CV grid search (Bi-GRU, CNN, Transformer) selects Bi-GRU as the best model. Its false-positive calls on *∼* 10 million held-out cytosines are harvested as hard negatives, redundancy-filtered and merged into an augmented training set, finally used to retrain and deploy transcriptome-wide our Bi-GRU model. **(c) Resources**. The resulting writer-specific prediction database, with refined motifs and coherent secondary-structure profiles are released for community use.

Next, we trained and compared three deep-learning architectures (Bi-GRU, CNN and a Transformer) in a multiclass setting to distinguish writer-specific methylation from unmodified cytosines. The results revealed performance ceilings: models struggled to separate genuine m^5^C sites from close lookalikes (Fig. 1b).

To address this, we used the best-performing architecture to mine false-positive predictions (i.e. hard negatives) from *∼*10 million unmodified cytosines withheld during pretraining. After augmenting the training set with these hard negatives, transcriptome-scale predictions recovered sharper writer motifs and the expected writer-specific secondary-structure patterns. To further validate model behavior, we assessed predictions across the whole transcriptome for functional enrichment, compared our models with state-of-the-art machine learning methods, and trained a surrogate model to interpret the models’ learned features and mutational effects. Finally, we released a high-confidence, writer-specific m^5^C catalog (Fig. 1c), together with trained models for community use.

### 2.1 Construction of a high-confidence catalog of m^5^C sites

Liu et al.^28^ processed the same dataset we decided to adopt^27^ by utilizing the iMVP clustering framework combined with multiple validation steps to assign m^5^C sites to four methyltransferase-specific classes: Type I/II/III/IV (corresponding to NSUN2/6/5/1). These enzymes represent the primary writers responsible for bulk mRNA 5-cytosine methylation in human cells (as further discussed in Suppl. Note 2).

However, manual inspection of the members of each cluster revealed weak boundaries and residual contamination by bisulfite artifacts. To quantify this, we used the motifs drawn from each NSUN cluster to score every candidate site against a background distribution (equation (1), Section 4.1.2). Notably, 15.1% of the sequences were more likely under the background model or matched the wrong cluster, particularly within the NSUN2 group (Type I). This prompted us to implement a more stringent clustering and filtering pipeline (Fig. 1a, Sect. 4.1).

First, to re-discover and refine writer-specific motifs, instead of clustering through sequence distance-base metrics as previously proposed,^28^ we utilized the STREME^29^ motif discovery algorithm. Indeed, motif discovery can adapt to the actual motif size, position, and entropy, easily filtering out sequences with no prevalent motif. In contrast, distance-based methods scale quadratically with the number of sequences thus forcing the use of heuristics with large datasets, such as fixing the window size uniformly across all sequences, reducing data dimensionality or using trivial distance measures. This may lead to the inclusion of uninformative flanks and disregard the entropy of each position and the different effects of substituting alternative bases. Such confounding factors can often dominate the distance measure, giving rise to fuzzy cluster boundaries rich in false-positive instances, especially with noisy data. On the other hand, as reported by Liu et al.,^28^ STREME stochastic initialization can lead to different outputs across runs, and cannot always recover all four clusters. To mitigate this issue, we repeated runs with different parameters and seeds, crucially modifying the objective function to “Central Distance.” Resulting motifs were then manually merged based on similarity and consistency with experimental evidence to build the final position-specific frequency matrices (PFMs). To recover STREME’s missed sites, these PFMs were used to rescore the entire dataset using Equation 1, and sites were retained only when the likelihood exceeded background (see Section 4.1 for details).

Figure 2a shows the final four motifs derived from these refined clusters. They closely match those reported in previous studies,^27, 28, 30–32^ but the Type I logo is notably sharper than in the original iMVP clusters, indicating that our filtering removed bisulfite-induced false positives that had disproportionately inflated the poorly defined Type I group.

**Figure 2:**
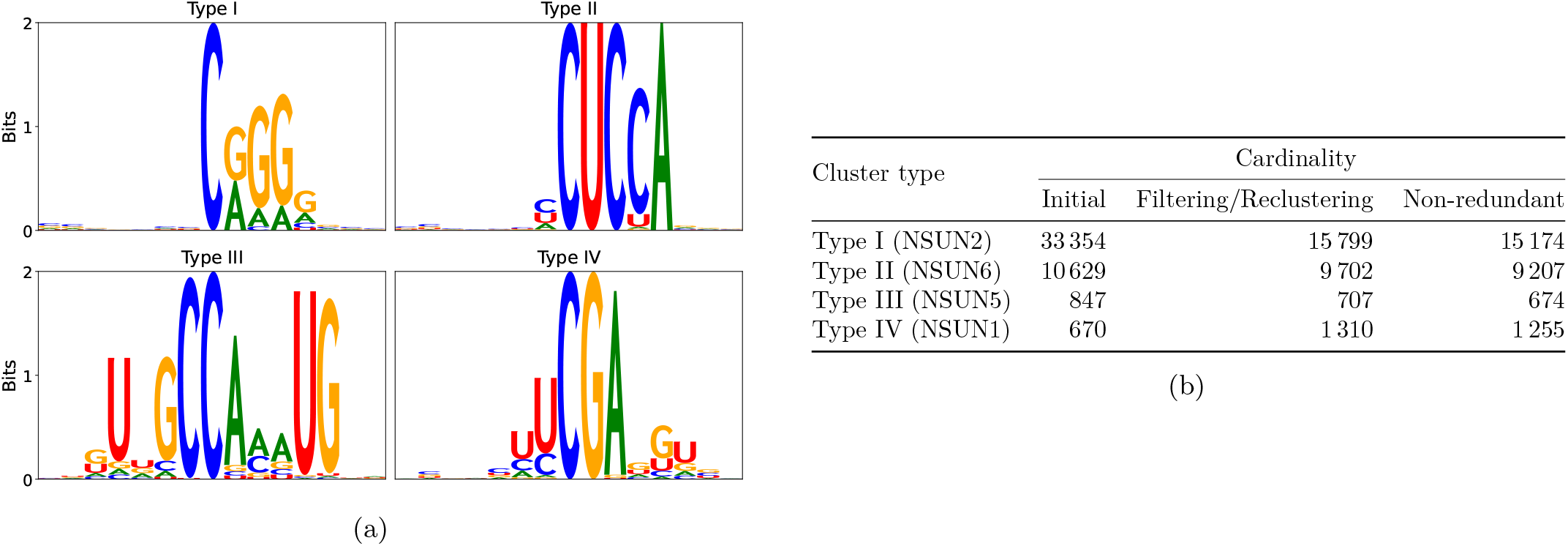
Refinement of writer-specific motifs and cluster sizes. (a) Final motifs for methyltransferase-specific sites (Types I–IV correspond to NSUN2, NSUN6, NSUN5, NSUN1). Letter sizes reflect information content (0–2 bits). (b) Cluster cardinalities before filtering (Initial), after filtering and re-clustering, and after redundancy removal.

To prevent data leakage, we then applied an exact custom script to remove sequences sharing more than 90% identity within the 21-nt motif core (Sect. 4.1.3). Unlike greedy approaches such as CD-HIT-EST,^33^ our non-greedy procedure ensures that similar sequences are consistently removed. Restricting comparisons to the conserved central core also avoids diluting similarity with unconserved distant flanks and provides a standardized basis for comparison with future algorithms that may use different window sizes.

Finally, we sampled an equal number of unmethylated cytosines from the same mature transcripts that harbored at least one modification. This strategy provides high confidence in their unmethylated status and guarantees that negatives derive from mature RNA, rather than untranscribed DNA, pre-mRNA or intergenic regions that would be trivially distinguishable from positives. The same redundancy filtering was applied to this negative set.

Overall, this process reduced the raw 45 500 candidates to a non-redundant set of 26 310 high-confidence sites distributed across the four principal methyltransferases, accompanied by an equal number of negatives. This curated catalog served as the foundation for all subsequent model training and evaluation. Sequence cardinalities after filtering and reclustering steps are summarized in Fig. 2b.

### 2.2 Convergence of the performance of our proposed deep models

We trained three modern architectures (Bidirectional GRU, Stacked 1D-CNN, and Transformer encoder) on our curated dataset to predict m^5^C modifications. Model architectures are described in Section 4.2.

Through exhaustive five-fold cross-validation and grid search on the training set (details in Sect. 4.3 and Suppl. Note 5), we evaluated several configurations for each architecture. Remarkably, the best configurations per backbone as well as across architectures showed minimal performance variation, with slightly lower metrics for the Stacked 1D-CNN. Average five-fold performance of the three best model per architecture are shown in Fig. 3a, with their hyperparameters indicated in Fig. 3b. Extensive Tables for the top 10 models as well as their hyperparameters are show in Suppl. Tables S1-S6.

**Figure 3:**
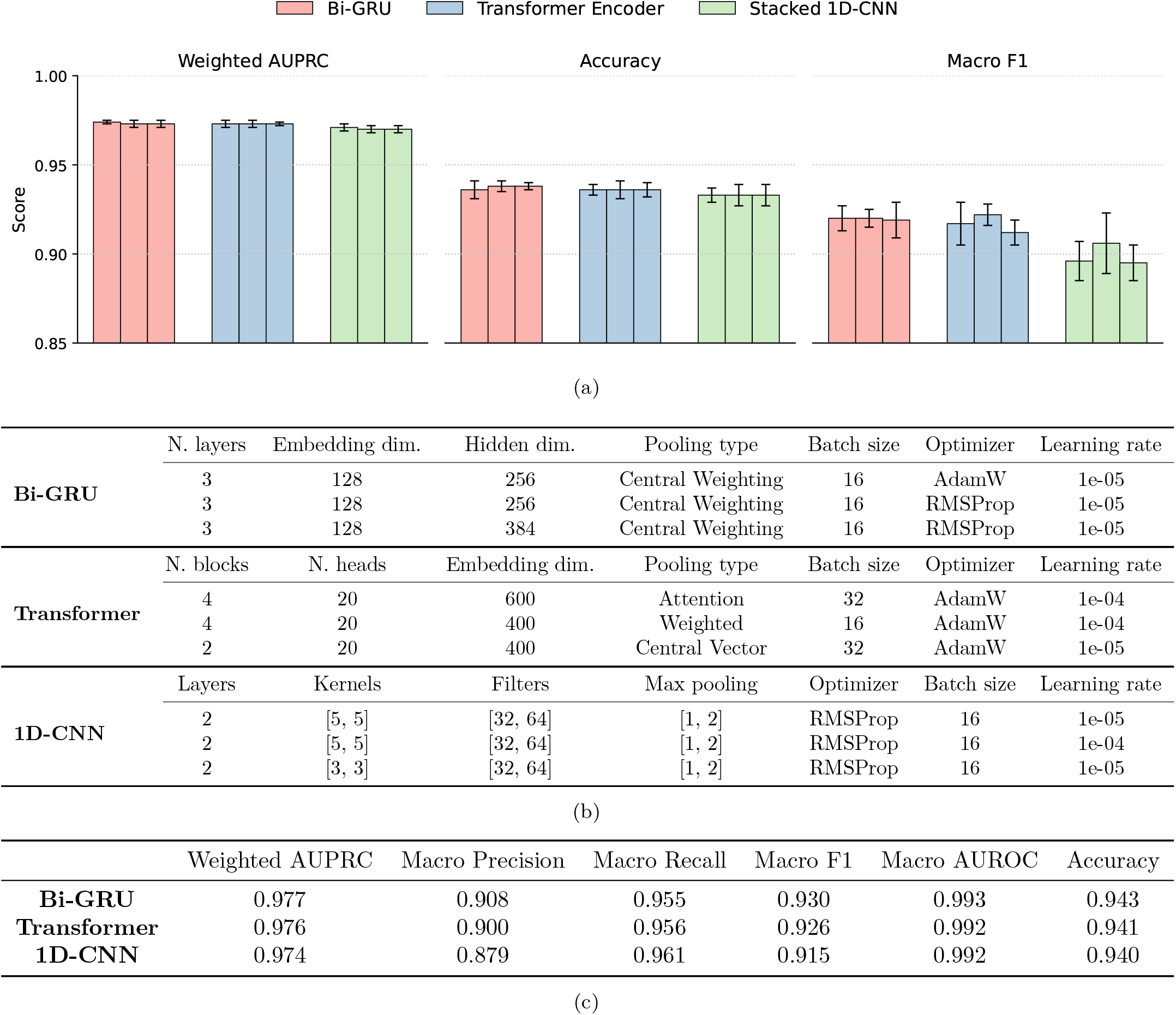
Best three models for each architecture (Bi-GRU, Transformer and 1D-CNN), obtained by grid search. (a) Average five-fold cross-validation performance on the training set of the three best models per architecture. (b) Hyperparameters of the same models. (c) Test set performance of the best configuration from each architecture after retraining on the full training set.

This convergence became even more pronounced when we retrained the best configuration from each architecture on the full training set (Fig. 3c). Even extending the sequence context provided no benefit: a Transformer grid search using 101-nt windows showed seven of the ten best models still originated from 51-nt configurations (Suppl. Table S7). The consistency across architectures and hyperparameters suggested that model capacity was no longer the main limiting factor. To understand the remaining errors, we analyzed confusion matrices generated by running each trained model on the complete dataset. As shown in Fig. 4, the Bi-GRU’s false positives predominantly contained sequences with motifs resembling those of genuine m^5^C sites. This pattern, replicated in the Transformer and CNN results (Suppl. Figs. S1-S2), revealed a critical insight: while our models effectively filtered out obvious negatives, they struggled to distinguish between true m^5^C sites and cytosines with similar but subtly different sequence contexts.

**Figure 4:**
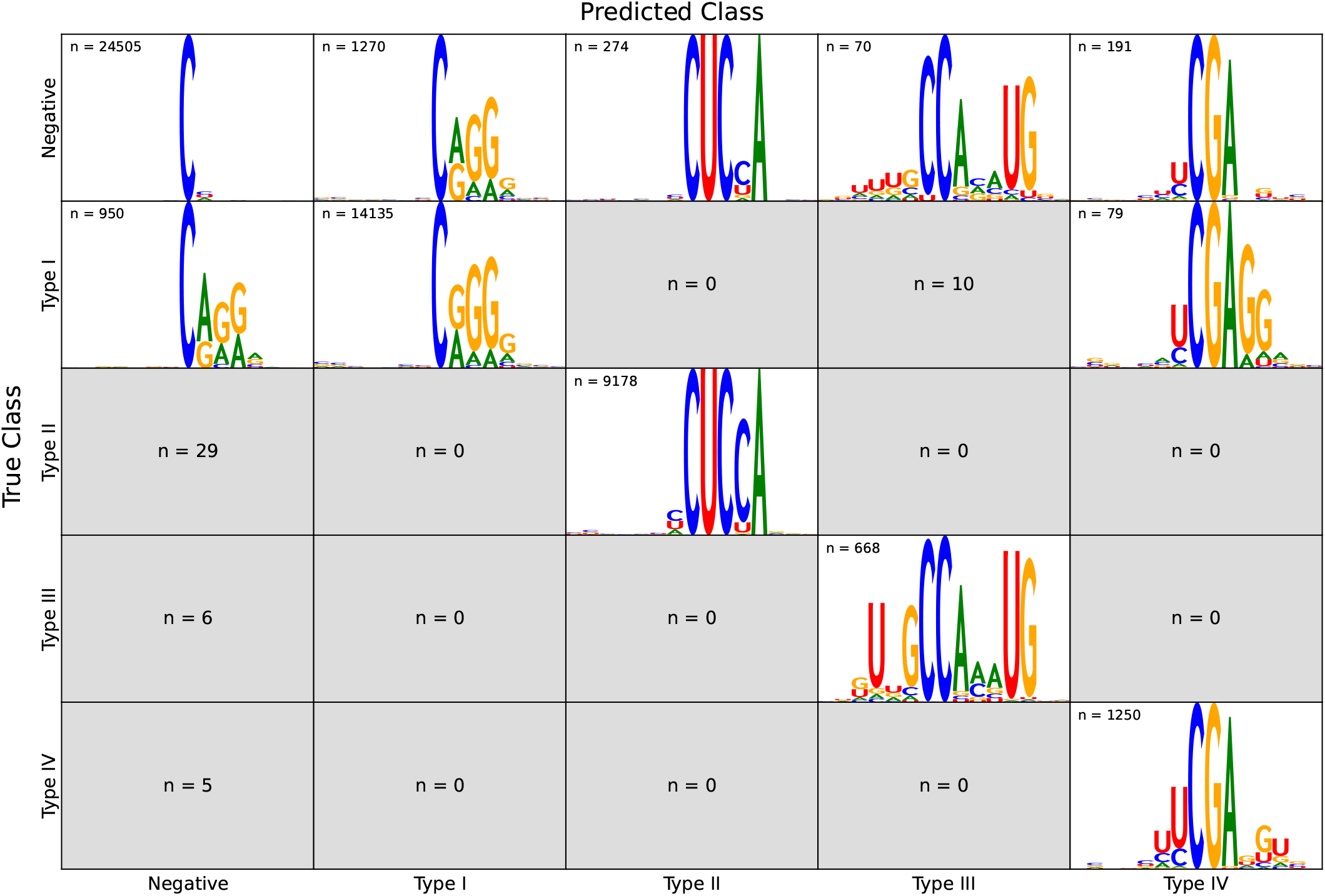
Performance of the final Bi-GRU on the combined training + test corpus. Rows are the true classes, columns the model’s predictions. Inside each cell we report: (i) the number of windows assigned to the (true,predicted) pair; and (ii) a sequence-logo summarizing the corresponding windows whenever *n* ≥ 50. Logos that would be based on fewer than 50 sequences are suppressed to avoid noisy motif representations.

### 2.3 Hard-negative retraining

This systematic misclassification of negatives with motif similarities to methylated sites indicated that our training set, despite careful curation, was not challenging enough. Standard negative sampling had primarily captured “easy” cases: cytosines with sequences clearly distinct from any writer motif. To address this limitation, we devised a targeted hard-negative mining strategy to individuate false m^5^C predictions whose motifs closely mimic those of true positive sites. This strategy loosely resembles boosting techniques,^34^ since it compels the model to learn from its own mispredictions and the associated biological context, effectively acting as a form of “biologically inspired” boosting.

#### 2.3.1 Light augmentation retraining

We implemented an initial hard-negative mining strategy using the *∼*10 million unmodified cytosines withheld from training. The best-performing Bi-GRU from each cross-validation fold was used to scan these cytosines to identify those misclassified as modified. After redundancy filtering, we augmented each fold’s training set with hard negatives sampled across different probability thresholds (see Methods 4.4).

Cross-validation performance showed modest improvements in weighted AUPRC, with better results when including hard negatives from higher probability bins (Suppl. Fig. S3). As expected, gains were limited because: (i) the unchanged validation sets contained predominantly easy negatives, and (ii) some high-confidence “negatives” may represent genuine methylation sites missed by bisulfite sequencing.

#### 2.3.2 Transcriptome-wide evaluation

To assess biological relevance beyond standard metrics, we applied both baseline and lightly augmented models to predict m^5^C sites across the transcriptome. We then evaluated the resulting predictions by examining RNA secondary structure with RNAfold^35^ and by assessing the sharpness and coherence of motifs for each methyltransferase-specific predicted site. Although the augmented model yielded clearer sequence motifs and more defined structural patterns compared to the baseline, its predictions still lacked robust signals (left and center columns, Fig. 5), suggesting over-reliance on trivial features.

**Figure 5:**
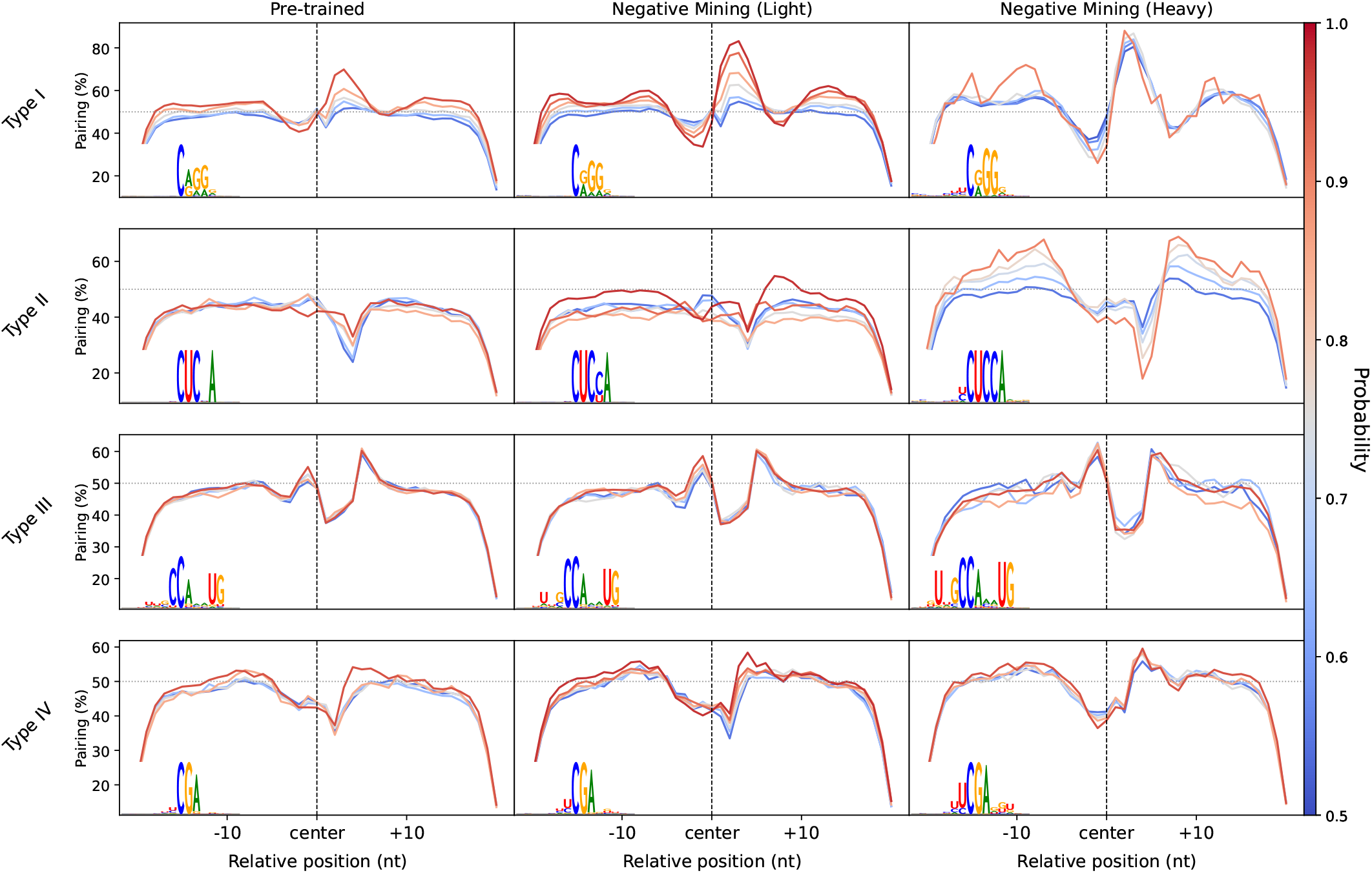
Predicted RNA secondary structure context improves as harder negatives augment training data. Each panel plots the percentage of nucleotides that are base-paired (y-axis) versus the distance from the central cytosine (x-axis). Predictions are performed with RNAfold. Rows correspond to the four NSUN classes; columns compare three model stages: (**left**) baseline Bi-GRU, (**center**) Bi-GRU after light hard-negative augmentation, (**right**) Bi-GRU after heavy hard-negative augmentation. Within every panel, colored curves group sites by oracle probability. Inset sequence logos summarize the motifs of the same site sets. The heavy-trained model not only sharpens writer motifs but also restores the writer-specific secondary-structure patterns reported experimentally.

#### 2.3.3 Heavy augmentation retraining

Building on the previous analyses, we systematically evaluated different hard-negative augmentation strategies by inspecting transcriptome-wide predictions of the trained models. The most substantial improvements arose from aggressively increasing the number of hard negatives in the training set and oversampling them within each mini-batch. The optimal framework is detailed in Section 4.4. Only after this augmentation did transcriptome-wide predictions recapitulate writer motifs with crisp base preferences *and* secondary-structure profiles consistent with experimental evidence (right column, Fig. 5):

- **Type I (NSUN2)**: Predictions now clustered at 5^*′*^ stem edges, matching published structural preferences^15^
- **Type II (NSUN6)**: Sites localized to hairpin loops with characteristic concave pairing profiles^30^
- **Types III and IV** – sequence logos sharpened and pairing scores improved modestly. The relatively limited structural gain may be attributable to the smaller training sets; however, writer-specific structural constraints for these enzymes have not yet been experimentally demonstrated, and the profiles we identified are consistent with the patterns reported by Liu et al.^28^

Together, these results highlight that saturating the training set with carefully selected hard negatives is essential for learning the subtle interplay between sequence motifs and RNA structure that distinguishes genuine m^5^C sites from spurious matches.

### 2.4 Gene enrichment analysis

Having confirmed that our final model learns both sequence and structural patterns characteristic of each writer, we next asked whether the predicted sites map to genes whose functions align with known methyltransferase biology. For each NSUN class, we selected transcripts harboring the highest-probability methylation sites (excluding any site used during training or testing), and queried `g:Profiler`^36^ against GO, KEGG, Reactome and the Human Phenotype Ontology (Methods, Sect. 4.5).

Figure 6a (GO Biological Process) and Figure 6b (KEGG pathways) show results for Type I sites (NSUN2), the most widely characterized methyltransferase, which accounts for most mRNA m^5^C sites. The top GO:BP terms: “regulation of neuron projection development”, “regulation of nervous system development”, “axonogenesis” and “forebrain development” directly link NSUN2 targets to neuronal growth and differentiation. These associations align with experimental data showing that NSUN2 loss accelerates neurodegeneration^37^ and that overexpression exacerbates astrocyte activation and Alzheimer pathology.^38, 39^ Consistently, biallelic NSUN2 deficiency causes developmental brain anomalies,^40^ and early neuroepithelial progenitors require NSUN2 for proper differentiation.^41^

**Figure 6:**
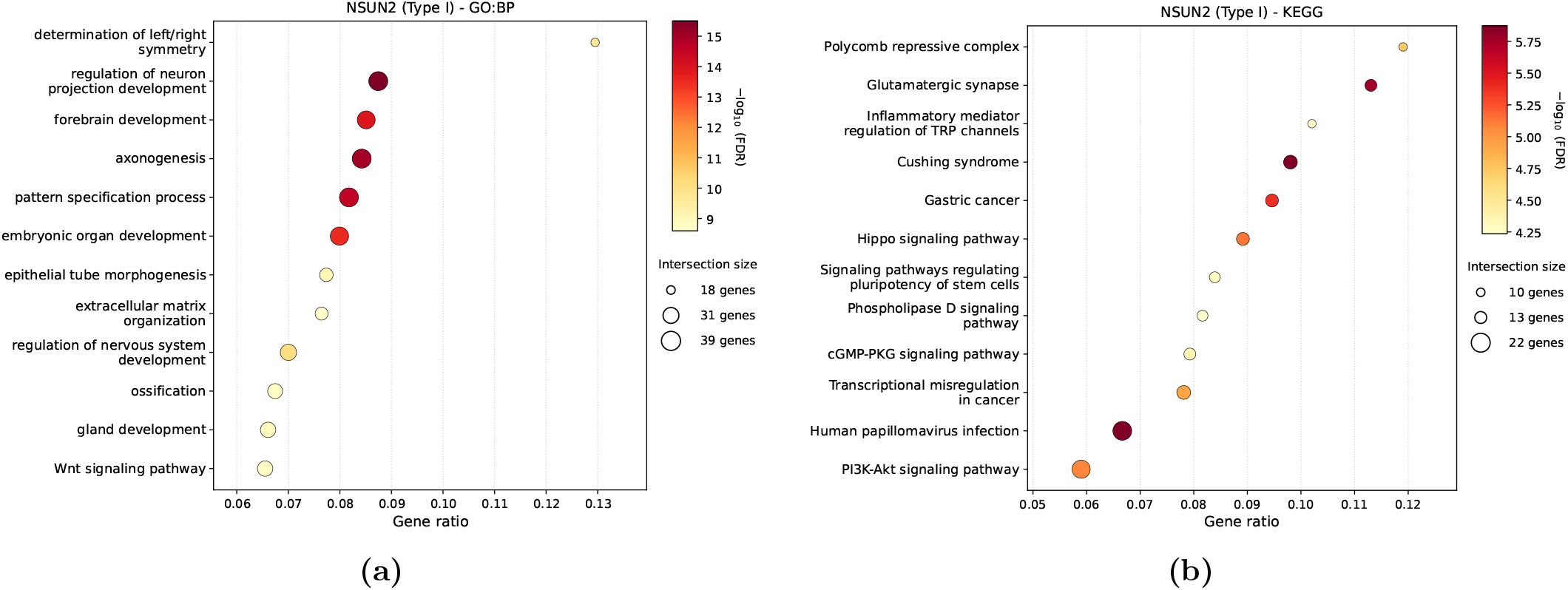
Functional coherence of NSUN2 (Type I) predictions. **(a)** Top Gene Ontology–Biological Process (GO:BP) terms enriched among genes harboring high-confidence NSUN2 sites. **(b)** Top KEGG pathways for the same gene set.

Additional developmental terms echo the m^5^C waves observed during embryogenesis.^27^ Both GO:BP and KEGG also highlight tumor-related pathways, including “extracellular matrix organization”, “Wnt signaling”, “PI3K-Akt signaling” and “transcriptional misregulation in cancer”, consistent with NSUN2’s role as an oncogenic driver.^42–45^

Furthermore, the other three writers showed equally compelling functional coherence (Suppl. Figs. S4-S7). For instance, NSUN5 involvement in gastric cancer and cardiac genes and NSUN6 methylated genes enriched for platelet activation, brain and embryonic development are also supported by experimental evidence.^27, 46–49^

Full enrichment tables are provided at Zenodo https://doi.org/10.5281/zenodo.16629378.

### 2.5 Comparison with state-of-the-art m^5^C predictors

To benchmark our approach against existing methods, we compared our best models with two m^5^C predictors: Deepm5c^24^ and MLm5c,^25^ that achieved better results with respect to other state-of-the-art approaches. Both Deepm5c and MLm5c leverage ensemble stacking techniques.^50^ Deepm5c includes as base learners both deep and traditional machine learning models and as meta-learner a 1D-CNN, while MLm5c uses only classical base learners such as Support Vector Machines and Random Forests and leverages a logistic-regression meta-classifier to detect m^5^C sites.

Since ours is the first multiclass m^5^C classifier, we converted our outputs to binary predictions (modified vs. unmodified) for fair comparison. It should be noted that both benchmark methods are trained on m^5^C sites from the m^6^A Atlas,^51^ with negatives sampled randomly from the genome, a critical difference from our transcript-matched approach. Training specifications and benchmark configurations are detailed in Suppl. Note 7.

#### 2.5.1 Testing models on our curated dataset and on m^6^A Atlas

On the Deepm5c test set, derived from the m^6^A Atlas, our models (without dataset-specific tuning and using only 80% of training data) matched or exceeded the complex ensemble’s performance: 86.2% accuracy (Transformer), 84.8% (Bi-GRU) and 84.7% (CNN) versus 85.2% for Deepm5c (Table S8). More surprisingly, only MLm5c’s simple composition-based classifiers dramatically outperformed both Deepm5c and our deep learning models on their dataset, with LightGBM achieving 91.5% accuracy using only trinucleotide frequencies.

This paradox, i.e. simple k-mer counts outperforming sophisticated deep learning, prompted us to examine the datasets more closely. When we trained the same LightGBM model on our data, its accuracy plummeted to 62.5% (training settings and implementations are described in Suppl. Note 7). The explanation lies in negative sample selection: our negatives come from the same transcribed mRNAs as positives, while theirs are drawn randomly from the genome, predominantly capturing untranscribed or intronic regions. A decisive experiment confirmed this: training LightGBM to distinguish our transcript-derived negatives from MLm5c’s genome-sampled negatives achieved 85.1% accuracy (Suppl. Note 7). The model learned to identify mRNAs versus genomic background, not m^5^C modifications. Consistent with this, Jensen–Shannon divergence of k-mer distributions (Fig. 7a) is greater between MLm5c negatives and our negatives than between our positives and negatives, explaining why composition alone suffices on their dataset but fails on ours.

**Figure 7:**
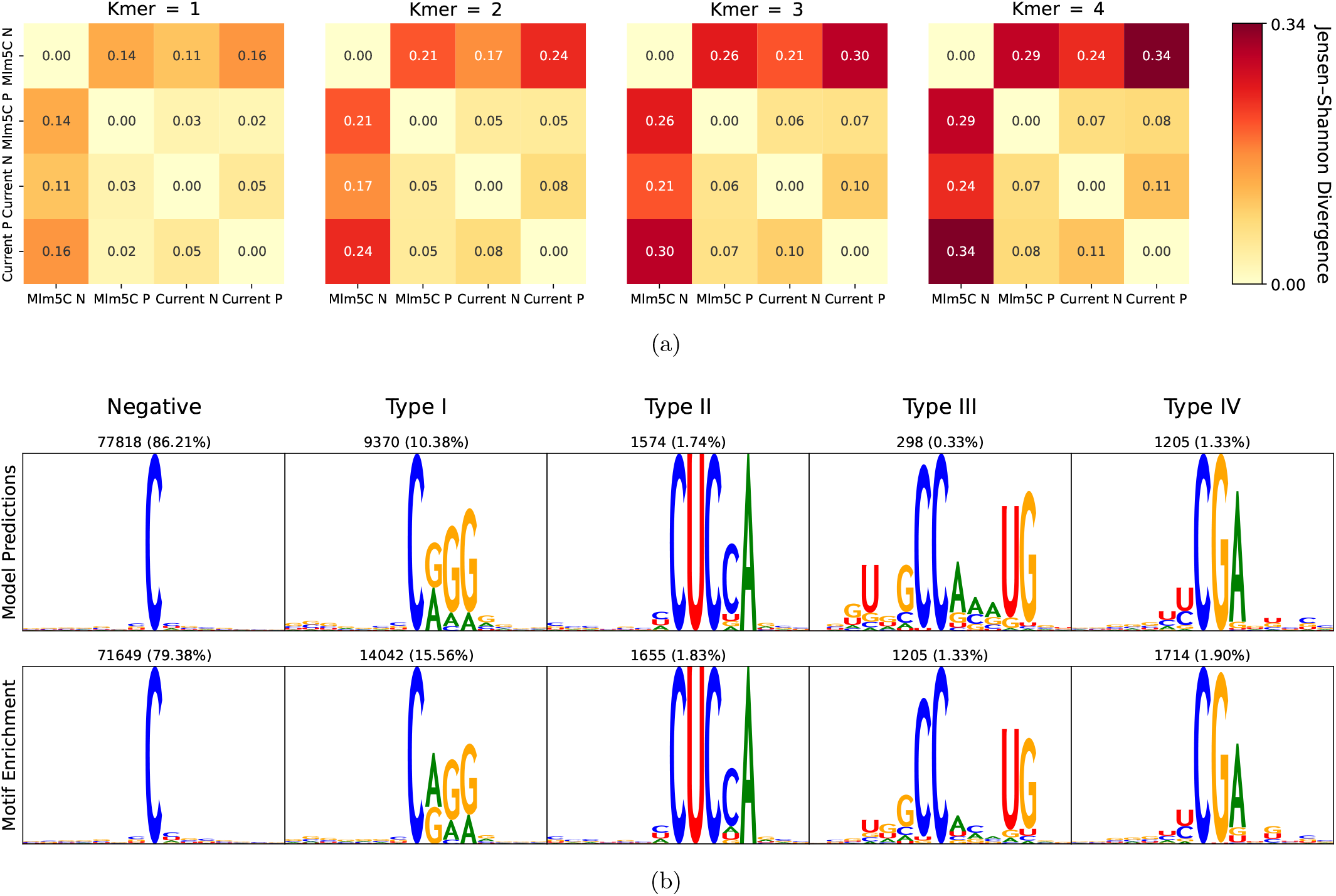
Comparison of our current and m^6^A datasets. (a) Confusion matrices of the Jensen-Shannon divergence between nucleotide k-mer distributions of MLm5c Positives (MLm5c P), MLm5c Negatives (MLm5c N), Current dataset Positives (Current P) and Current dataset Negatives (Current N). (b) Top row shows the motifs drawn from the predictions of the Transformer model on the m^5^C sequences of the m^6^A Atlas. The bottom row depicts the motifs drawn from the same dataset, but through motif enrichment by utilizing the likelihood of the position frequency matrices of the four NSUNs proteins through equation (1).

#### 2.5.2 Atlas dataset contains predominantly spurious m^5^C sites

Despite the apparent simplicity of distinguishing genomic background from mRNA sequences, our models achieved markedly lower performance on the m^6^A Atlas dataset compared to our curated data. This discrepancy prompted us to investigate the quality of the Atlas m^5^C annotations themselves. To investigate, we applied our Transformer (trained on our dataset) together with motif log-likelihood scoring (equation (1)) to the Atlas m^5^C sequences (Suppl. Note 7). This orthogonal validation method provided an independent check on site authenticity.

The results were striking: 86.21% of Atlas sites were classified as unmodified by the Transformer, while 79.38% scored below the motif likelihood threshold (Fig. 7b). Moreover, direct STREME runs on the Atlas failed to recover any of the four canonical NSUN motifs, indicating that the Atlas contains predominantly spurious m^5^C calls.

Notably, we selected the base Transformer (without hard-negative retraining) because it exhibited the most lenient classification behavior, particularly for Type I (NSUN2) sites. This choice ensured that even marginal motif-like sequences with suboptimal secondary structure features would still be classified as positives, providing the most generous possible assessment of the Atlas sites. Furthermore, PFMs derived from sites predicted as positive by both motif log-likelihood and the Transformer closely match biochemically characterized motifs for NSUN1, NSUN2, NSUN5, and NSUN6,^27, 28, 30–32^ reinforcing that negatively predicted sites do not resemble known NSUN motifs.

We considered whether other human m^5^C writers might account for the remaining Atlas sites. However, the known alternative methyltransferases operate in restricted contexts: NSUN3 and NSUN4 are confined to mitochondria,^52^ while NSUN7 specifically targets enhancer RNA^53^ (detailed in Suppl. Note 2), and these specialized writers cannot plausibly explain the > 75% false-positive rate observed across the Atlas.

### 2.6 A surrogate model explains the m^5^C predictions

We next sought to interpret what the final Bi-GRU actually learned about how single-nucleotide changes affect motif-specific cytosine 5-methylation. To this end, following Seitz et al.,^54^ we built an *in silico* multiplex assay of variant effects (MAVE) by systematically mutating transcriptome-wide sites predicted as methylated and rescoring each sequence with the trained Bi-GRU (the “oracle”). This yields a dataset of natural and mutated windows paired with oracle-predicted methylation probabilities.

To make these effects interpretable, we fit methyltransferase-specific *surrogate models* with MAVE–NN^55^ (details in Suppl. Notes 8 and Methods Sect. 4.6). A surrogate is a deliberately simpler model trained to imitate the oracle but whose parameters can be directly interpreted. MAVE–NN estimates the linear contribution of each nucleotide at each position and their second order epistatic interactions. Large positive values of the linear coefficients raise the predicted probability of cytosine modifications, while negative values lowers them. Second order coefficients quantify the departure from additivity: large positive (negative) values indicate synergistic (antagonistic) epistasis between the two substitutions. Therefore, the model is able to learn the mutational effect of each nucleotide at each position of a given motif as well as potential epistatic effects (Suppl. Note 9). Together, these coefficients provide a quantitative, position- and pair-level map of the variant effects the Bi-GRU has internalized, which we analyze below by class.

We first verified that the surrogates faithfully approximate the oracle. On held-out test sequences, surrogate predictions tracked the oracle across all four NSUN classes (Fig. 8a), with the best fits for the abundant classes (Type I and II) and lower ℛ ^2^ for the sparse classes. This pattern is expected: a smaller dataset provides fewer positive windows available to seed the *in silico* MAVE, yielding sparser coverage of sequence space and a narrower dynamic range of oracle scores.

**Figure 8:**
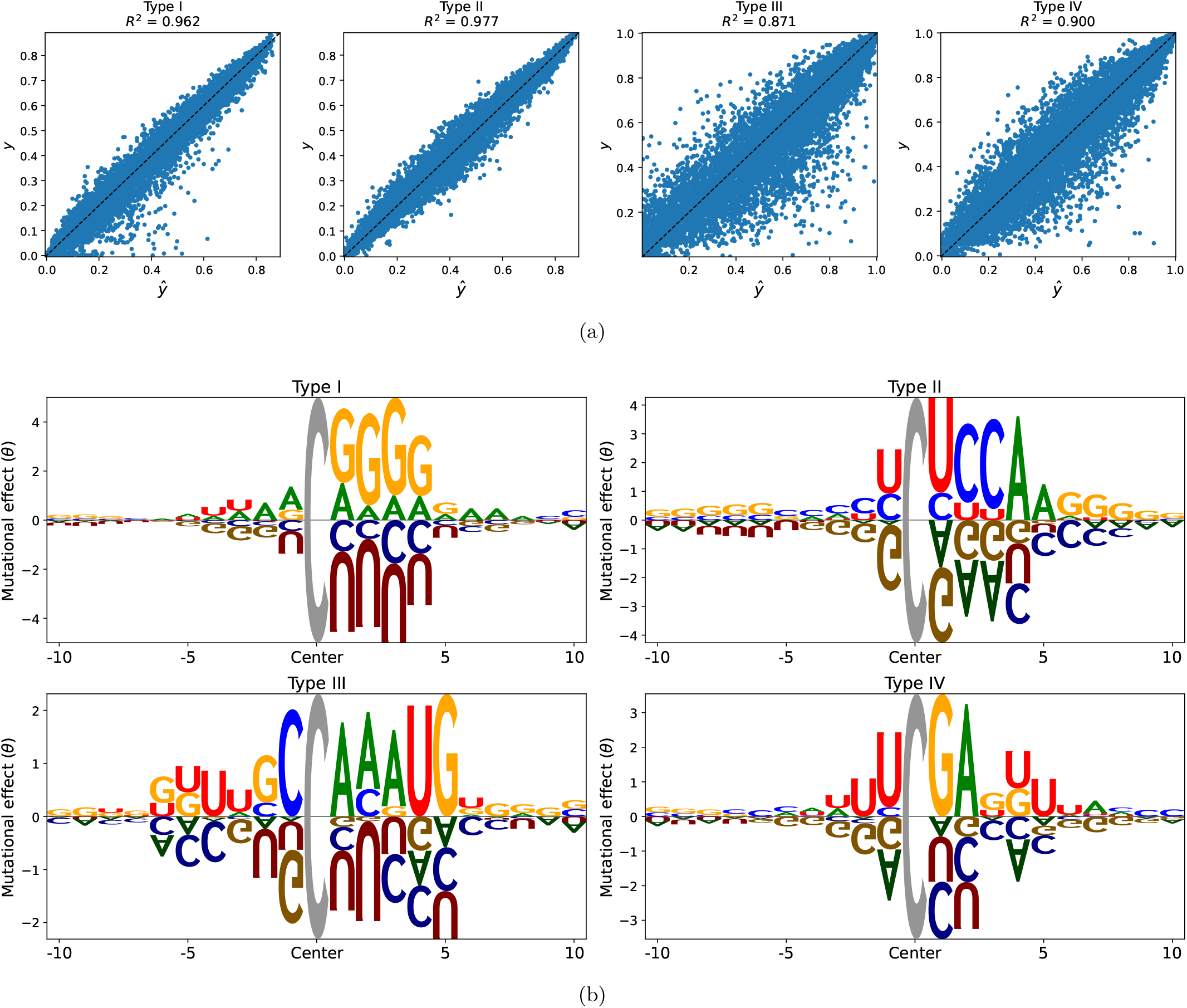
Surrogate model analysis. (a) Correlation between Surrogate Model predictions (*ŷ*) and oracle outputs (*y*) for the four positive classes on the test set. Coefficients of determination are also reported. (b) Additive effects captured by the surrogate model linear parameters *θ*s trained with the mutational dataset constructed from each m^5^C positive class. Large positive *θ* values raise the predicted modification probability, whereas large negative values lower it.

With surrogate fidelity established, we turn to interpretation. Across all classes, positive additive coefficients recapitulate the canonical NSUN motifs (Fig. 8b), whereas large negative values for alternative bases highlight disruptive effects learned by the oracle. Unlike consensus PFMs, which average over sequences already predicted as methylated, the additive maps expose position-specific effects that such averages can hide. A clear example is Type II: its additive coefficients show positive contributions from pyrimidines downstream of the modified cytosine, an asymmetry absent from the consensus PFM but plausibly reflecting steric constraints on NSUN6 binding. For the same class, mildly positive coefficients for guanines and cytosines outside the core CUCCA motif are consistent with higher base-pairing and stacking energies that may stabilize the hairpin surrounding the NSUN6 binding site. For Type I (NSUN2), we instead observe a local enrichment of purines with a depletion of U/C immediately downstream of m^5^C, consistent with stronger same-strand stacking that stabilizes the stem edge during the cytosine-flipping step required for methylation. Full-length additive parameters and pairwise interaction maps are shown in Suppl. Figs. S8-S9. The interaction heatmaps reveal pronounced clusters of strong epistasis in the immediate vicinity of the motif core, indicating that combinations of substitutions at these positions have non-additive effects on methylation probability.

## 3 Discussion

Chemical modifications add a regulatory layer to RNA that affects splicing, nuclear export, localization, stability, and translation, with widespread links to development and disease. Comprehensive, accurate maps of these marks are therefore essential for understanding both biology and medicine. To this end, several experimental protocols have been developed to locate modified sites transcriptome-wide. However, all introduce characteristic biases, false positives arising from protocol artifacts and systematic false negatives where certain sequence or structural contexts are under-detected, leaving current catalogs often noisy and incomplete. These limitations have motivated machine learning algorithms to identify experimentally missed sites. However, an architecture-first focus (without rigorous dataset design and curation) means models are trained on partial, noise-contaminated labels and therefore propagate those artifacts, producing misleading gains on biased benchmarks. To make matters worse, relying solely on these benchmarks (without orthogonal, biology-driven validation) can conceal model failure.

m^5^C provides a clear case study. Several recent models report impressive scores, but much of the results traces to label-design biases: datasets composed of “negatives” sampled at random from genomic DNA and “positives” coming from mature mRNAs make simple nucleotide composition highly discriminative. Furthermore, current m^5^C datasets compound this problem with widespread false positives: over 75% of the sites in the m^6^A Atlas lack any trace of known m^5^C writer motifs. The design of a biology-driven analysis (motif concordance to known writers and k-mer composition) revealed that the apparent gains largely reflected dataset artifacts rather than genuine m^5^C signal.

Therefore, we decided to invert the paradigm. First, we reconstructed a high-confidence writer-resolved catalog of 26 310 human m^5^C sites from the extensive developmental dataset Liu et al.^27^ We reduced the noise that has plagued previous efforts by applying rigorous quality control, including motif-guided filtering, writer-specific clustering, and exact redundancy removal. Differently from recent works, negative cytosines were drawn from mature transcripts bearing at least one modified site, providing the highest confidence in their unmodified status while also minimizing nucleotide-composition bias.

Next, we designed three architecturally distinct models (Bi-GRU, CNN, and Transformer) that predict m^5^C sites and also distinguish the methyltransferase classes underlying these modifications. To our knowledge, this is the first multiclass model for m^5^C prediction. When we trained these different models on our curated dataset, all converged to nearly optimal performance (AUPRC > 0.97). A biological analysis of motifs among the few negatives misclassified as positives revealed why residual errors persisted: miscalls clustered in unmodified cytosines whose local sequence closely resembled m^5^C contexts. Even with unbiased, transcript-matched negatives, the bottleneck was not model capacity but the scarcity of sufficiently challenging examples. Had we relied on metrics alone, this problem would have gone unnoticed. We addressed this by mining hard negatives from *∼* 10 million unmodified cytosines withheld during pretraining. By augmenting the training set with such look-alike decoys did we force the models to learn discriminative features beyond simple patterns. The impact was clear: sequence motifs among positives sharpened to match biochemical specificity, and the models began to leverage RNA structural constraints that were never explicitly programmed, including writer-specific secondary structure patterns experimentally reported: NSUN2 sites clustering at stem edges,^15^ NSUN6 preferring hairpin loops.^30^

To interpret what the final predictor actually learned (not just how well it scores) we fit class-specific surrogate models to an *in silico* multiplex assay of variant effects, using our model as the oracle. Analysis of the surrogate parameters revealed that our model learned the NSUN motifs across all four classes and quantified the effects of substitutions that disrupt methylation. Given that m^5^C dysregulation is implicated in multiple diseases, these results underscore the potential use of our model to prioritize variants that may impair m^5^C and contribute to disease risk. Furthermore, Transcriptome-wide predictions for all four writers were enriched in pathways matching their known functions. The concordance between computational predictions and experimental biology validates our approach and suggests that predicted sites represent *bona fide* methylation targets.

By releasing our AI models, curated training data and predicted sites at transcriptome-wide level, we aim to accelerate the transition from computational inference to biological discovery.

Summarizing, our results show that once model capacity is adequate, incorporation of domain knowledge through targeted data curation and biological validation matters more than model sophistication. Genuine gains stem from harder, better-curated data, not only from ever more elaborate architectures, which has been the prevailing emphasis of RNA-modification predictors to date. In perspective, we envision this pipeline to be extended to other RNA modifications, as only through the integration of computational and experimental rigor we can build tools that truly advance our understanding of the epitranscriptome.

## 4 Materials and methods

### 4.1 Construction of a high-confidence catalog of m^5^C sites

Through their iMVP clustering approach, Liu et al.^28^ were able to draw and validate the motif of the sequences within each cluster associated to the four methyltransferases NSUN2, NSUN6, NSUN5 and NSUN1. However, manual inspection of the four clusters revealed sequences having windows with flanks missing the expected motif. Therefore, despite the more stringent filtering, and considering the still high false positive rate of BS-seq thoroughly discussed in Suppl. Note 1, we hypothesized that false positive sites were still present within the dataset. To overcome this problem, we designed a more stringent pipeline for false positive removal and motif clustering.

#### 4.1.1 Transcriptome mapping of human m^5^C sites

Starting from the 45 500 human m^5^C sites reported by Liu et al.,^27^ sites were mapped onto the hg19 reference genome, assigned to GENCODE v19 transcripts,^56^ and restricted to mature, poly(A),+ exons captured by the bisulfite-sequencing protocol. Additional filtering steps are detailed in Suppl. Note 3.1.

#### 4.1.2 Writer-specific motif reclustering using a modified STREME algorithm

To analyze iMVP^28^ clusters, we built position frequency matrices (PFMs) using the member sequences of each cluster. We then scored every site against each PFM by computing the average log-likelihood ratio:

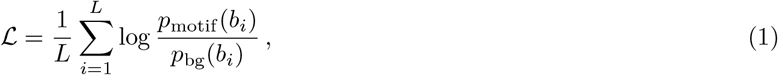

where *p*_motif_(*b*_*i*_) is the PFM probability of base *b*_*i*_ at position *i*, and *p*_bg_(*b*_*i*_) is a position-matched background frequency estimated from 100 000 unmodified cytosines in the same transcripts. The motif length *L* was defined by retaining only low-entropy flanking positions. For NSUN6 (Type II), for example, we used positions [− 1, …, +4], since more distant columns showed no bias. A positive ℒ indicates the sequence is more likely under the motif model than background; a negative ℒ indicates the opposite. Interestingly, we found 15.1% of the sequences were either more likely under the background distribution or under the wrong cluster. This prompted us to recluster the methylated sites. To overcome the limitations of previously proposed sequence distance-based clustering algorithms^28^ (Section 2.1), we utilized the STREME^29^ motif discovery algorithm.

However, STREME’s stochastic initialization can sometimes lead to slightly different or even missing motifs. Therefore, we reran STREME nine times (with 3 different seeds x 3 different length parameters) by including the methylated cytosines in the center and the respective 25-nt flanks. Since motif lengths can vary, we set the maximum motif length to 8, 9, or 10 nt for each seed. Minimum length was kept at 5. The central cytosine position allowed us to change STREME objective function to “Central Distance”, which drastically improved the results. Finally, we manually merged clusters from the nine runs based on motif similarity with experimental evidence. Rationale on the choice of the algorithm, the advantage in using central distance over traditional distance metrics, and further details are provided in Suppl. Note 3.2.

To recover any true sites missed by STREME, we rebuilt PFMs from the refined clusters and rescored every window using equation (1). Each site was retained and labeled by its highest-scoring motif only if that motif’s score satisfied ℒ > 0. This step removes windows lacking any motif signal, most of which represent BS-seq false positives. Sequence motifs throughout this work were drawn and visualized using `Logomaker`.^57^

#### 4.1.3 Redundant m^5^C sites filtering

To prevent data leakage between training, validation, and test splits, we first removed highly similar sequences from the dataset. A survey of the literature shows that most RNA modification detection pipelines use `CD-HIT-EST`^33^ to cluster sequences above a user-defined identity threshold; however, this greedy algorithm can assign very similar sequences to different clusters. Instead, we implemented a custom script that retains a candidate sequence only if its identity with *every* previously kept sequence falls below the threshold; otherwise, the candidate is discarded, which also avoids the construction of a memory-constrained distance matrix.

A further complication is that apparent sequence similarity depends on the length of the window used for comparison. Close to an m^5^C site the few nucleotides that form the writer-recognition motif are usually conserved at the *primary-sequence* level, whereas the more distant bases at most serve to establish the *secondary structure* (base-pairing pattern).^58^ Because the same pairing pattern can be realised by many different base choices (e.g. compensatory mutations in paired positions), secondary structure tolerates far more variation than primary sequence. Two windows can therefore look almost identical at their centers yet differ markedly in the flanks, so computing identity over the full length would dilute genuinely redundant cores with irrelevant peripheral mismatches and thus *underestimate* redundancy where it matters most. To focus on the informative region, we limited identity calculations to the central 21 nt (10 nt upstream, the modified cytosine, and 10 nt downstream) and applied a 90% identity cutoff. This method also ensures that redundancy filtering is not influenced by the length of the flanks considered around the modified site, which is often different among RNA modification detection algorithms.

#### 4.1.4 Negative Sites Selection

An equal number of non-redundant negative sites were drawn at random from a pool of *∼*10 million cytosines in the same mature transcripts that harbored at least one modification, ensuring these negatives were highly likely to be unmodified. The redundancy filtering procedure for negatives corresponds to the one used for positives.

### 4.2 Model Architectures

We trained three modern sequence-learning architectures: recurrent neural network (Bidirectional Gated Recurrent Unit – Bi-GRU), one-dimensional Convolutional Neural Network (1D-CNN), and Transformer encoder to predict m^5^C in a five-class setting. In particular, each model receives a window centered on the candidate cytosine and decides whether that cytosine is unmodified or modified by one of the four methyltransferases.

#### 4.2.1 Sequence encoding

All models ingest a fixed-length window centered on the candidate cytosine. If a cytosine is close to the sequence start/end, we left- or right-pad the missing flank with the dummy base P. Padding positions are encoded as zero vectors and masked out where appropriate (e.g. during Transformer attention or pooling).

Except for the ENAC representation used by the stacked 1-D CNN (Section 4.2.3), all nucleotides are treated as single tokens. Alternative schemes such as di- or tri-nucleotide tokens were pilot-tested but yielded no performance gain.

#### 4.2.2 Bidirectional GRU

First, each nucleotide in the sequence is converted to an embedding vector 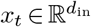, where *t* is the position in the sequence. Such embedded sequence is fed to a Bidirectional Gated Recurrent Unit (Bi-GRU).^59^ The bidirectional scheme sweeps the sequence in both the 5^*′*^→ 3^*′*^ and 3^*′*^→ 5^*′*^ directions, capturing context that precedes and follows every nucleotide, while the GRU cell itself offers a lighter, faster alternative to an LSTM. A summary of one GRU layer at time-step *t* is reproduced below (batch index omitted for clarity):

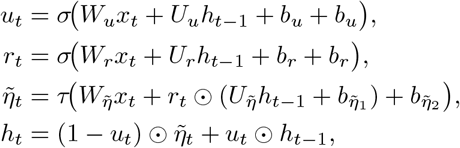

where 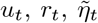 and *h*_*t*_ denote the *update gate, reset gate, new gate* (hidden candidate state) and *hidden state*, respectively. 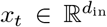 is the input embedding at time *t*. The parameters are weight matrices 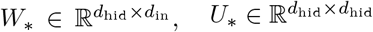, and bias vectors 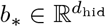. *σ* is the sigmoid function, *τ* is the hyperbolic–tangent, and indicates the Hadamard (element-wise) product.

The concatenated output 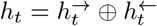 of the forward and backward GRUs forms a context matrix:

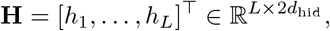

*L* being the sequence length. Different pooling architectures (described in Suppl. Note 4.1) have been explored to convert *H* to a context vector *c* passed through a fully connected (FC) layer (described in Suppl. Note 4.2) to make the multiclass classification.

#### 4.2.3 Stacked 1D-CNN

To extract spatial features from RNA windows we employ a stacked one-dimensional Convolutional Neural Network (1D-CNN); each Conv1D layer is followed by a Rectified Linear Unit (ReLU) non-linearity, dropout and a max-pooling layer.

##### Input encoding

The nucleotide string is converted to a two–dimensional tensor 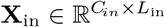, where the first axis enumerates channels and the second axis enumerates sequence positions.

- **One-hot**. Each nucleotide {A, C, G, U} is mapped to a four-component one-hot vector; hence *C*_*in*_ = 4.
- **ENAC** (enhanced nucleotide composition). A sliding window of size *k* scans the sequence of length *L*_seq_ (stride = 1). At position *i* the four channels hold the counts of {A, C, G, U} observed inside the window that starts at *i* and ends at *i* + *k* − 1. The resulting length is *L*_in_ = *L*_seq_ − *k* + 1, but the channel count is still four.

The subsequent CNN treats the encoded tensor identically regardless of the chosen representation.

##### Convolution

Given 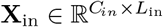, a one-dimensional convolution with *C*_*out*_ kernels of width *D* (stride *S* = 1, no dilation) produces the outputs:

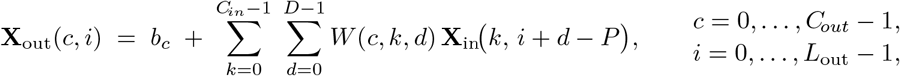

where 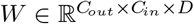 are the weights and 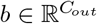 the biases. With padding *P* on both sides, the output length is

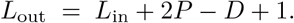

##### Activation and Dropout

Each element of **X**_out_ is passed through ReLU(*x*) = max(0, *x*), and dropout is applied after the non-linear activation function.

##### Max-pooling

One-dimensional max pooling is applied along the length axis. Let the ReLU output be 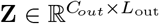.

With a pooling window of width *p* and stride *s* the operation is

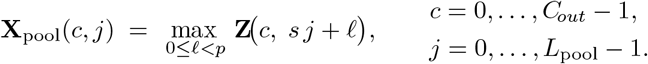

We set *p* = *s* (non-overlapping windows), giving *L*_pool_ = *L*_out_*/p*. The channel dimension remains *C*_*out*_ because pooling is performed independently on each filter’s activation map.

##### Network stack

The sequence [Conv → ReLU → Dropout → MaxPool] is repeated *n* times. In our experiments, we tested *n* ∈ {2, 4} . The final feature map 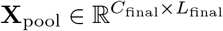 is flattened and sent to a fully-connected layer that yields the representation used by the classifier (Suppl. Note 4.2).

#### 4.2.4 Transformer encoder

We selected an encoder-based Transformer model, leveraging its self-attention mechanism to capture both short- and long-range interactions between nucleotides in the RNA sequence (an essential capability for our task given the presence of structural and motif-dependent constraints).

Each nucleotide in the sequence is converted to an embedding vector 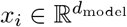, where *i* is the position in the sequence of length *L*. The resulting embedding matrix *X* = [*x*_1_, …, *x*_*L*_]^T^ ∈ R^*L×d_* model^ encodes the overall RNA sequence. In our experiments, we tested *d*_model_ ∈ {400, 600}.

##### Overall Structure

A stack of *n* encoder blocks then refines *X*. Each block consists of

1. **Layer normalization**. The input is normalised to zero mean and unit variance in the feature dimension.
2. **Query, Key and Value**. From the normalised input 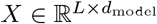 we obtain:

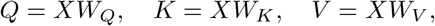

Where 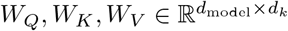.

3 **RoPE**. Rotary positional encodings (RoPE^60^) are applied to the rows of *Q* and *K*, injecting both absolute and relative positional information while preserving the dot-product form required by attention.
4 **Multi-head self-attention**. For each head *i* = 1, …, *h* we compute:

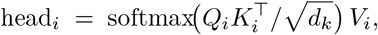

Where 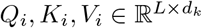. The heads are concatenated and projected back to the model width:

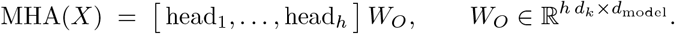

Multiple heads let the block attend to different patterns simultaneously.

5 **Residual connection**. The attention output is added to the block input.
6 **Layer normalization** (second instance).
7 **Feed-forward network**. A two-layer linear network with the SwiGLU activation^61^ is applied position-wise.
8 **Residual connection**. The FFN output is added to the block input, completing the Transformer block.

##### Stack and pooling

We stack *n* such blocks to obtain the output hidden matrix 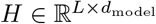. In our experiments, we tested *n* ∈ {2, 4}. Six pooling strategies described in Suppl. Note 4.1 are applied to map *H* to a context vector 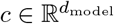 subsequently processed by the shared classification head (Suppl. Note 4.2).

### 4.3 Model training

The full dataset was split in a stratified manner, with 85% reserved for training and 15% held out as an independent test set.

Within the training partition, we performed five-fold cross-validation to select the best hyperparameter configuration for each of the three architectures described above. Stratification was maintained across folds. During optimization, mini-batches were sampled with a 50% probability of drawing a negative, while the remaining probability was equally divided among the four positive classes to counteract class imbalance.

An exhaustive grid search explored 360 Bi-GRU, 320 CNN, and 192 Transformer configurations. All windows were 51-nt long (cytosine at position 26); for the Transformer half of the configurations were also tested on 101-nt windows to assess the benefit of a longer context, as it is the most suited architecture for such task. The sequences were trained with early stopping on the validation set based on a weighted AUPRC. Additional information regarding grid search methodology and hyperparameter selection is provided in Suppl. Note 5.

#### 4.3.1 Evaluation metrics

We reported micro, macro and weighted AUPRC, AUROC, F1-score, precision, recall and accuracy for all the considered prediction tasks.

Plain micro- or nacro-AUPRC can be skewed: the former is dominated by the majority class, whereas the latter treats each class equally, making a single positive class as influential as the negative class. To balance these extremes we assign half of the total weight to the negative class and distribute the other half evenly across the four positives:

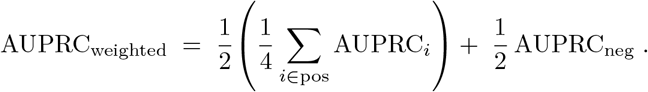

#### 4.3.2 Implementation

All experiments were run on NVIDIA A100-80 GB GPUs under PyTorch 2.1.0. Training employed `torch.cuda.amp.autocast`, enabling *automatic mixed precision*: numerically safe ops run in FP16/BF16 while the remainder stay in FP32, reducing memory use and training time with no loss of accuracy. For the Transformer models we replaced the standard attention kernel with `FlashAttention-2`,^62^ yielding an additional speed-up.

### 4.4 Hard-negative retraining

To improve model performance on challenging cases, and in particular for negative m^5^C sites wrongly predicted as positives, we implemented a two-stage hard-negative mining strategy using the *∼*10 million unmodified cytosines withheld from initial training.

#### 4.4.1 Hard negatives identification

We applied the best-performing architecture (Bi-GRU) from our five-fold cross-validation to scan all withheld cytosines. Any unmodified cytosine misclassified as modified was tagged as a “hard negative”. To prevent data leakage, these candidates were filtered using the same redundancy criteria described in Section 4.1, both against one another and against the initial dataset.

#### 4.4.2 Light augmentation

Given our Bi-GRU models trained on each of the five cross-validation folds, we:

- Loaded the trained model checkpoint
- Scanned the withheld negatives to identify hard cases
- Sampled uniformly hard negatives using six different probability upper bound ranges: [0.2–0.5], [0.2–0.6], …, [0.2–1.0] with 0.05 bin sizes
- Added *N*_0_*/*4 hard negatives per NSUN class (totaling *N*_0_ new negatives, where *N*_0_ is the original negative set size)
- Retrained from the saved checkpoint while maintaining the original 50:50 negative:positive mini-batch sampling ratio

Model comparison was based on weighted AUPRC averaged across the five folds. Crucially, validation sets remained unchanged from the original splits to avoid biasing our evaluation toward hard-negative performance.

Since the best model was obtained using hard negatives with a probability upper bound of 1.0, we repeated the augmentation procedure with this setting on the full training set to derive the final model.

It is worth noting that we did not limit sampling to just the highest-probability bins, but to all of them, to reduce the risk of incorporating too many potential false negatives that may have been missed by BS sequencing.

#### 4.4.3 Heavy augmentation

To enable a more biologically aware model comparison, we analyzed transcriptome-wide m^5^C predictions from both the pretrained and the hard-negative–augmented models (as described in Sect. 4.5.1). We examined the predicted sites in terms of their sequence motifs and secondary structures; the latter were folded with RNAfold from the ViennaRNA Package^35^ at 37°C (*Homo sapiens*).

Despite motif refinement and stronger secondary-structure signals in the lightly hard-negative–augmented model, we still observed insufficient structural features and weak motifs. Therefore, we tested more aggressive hard-negative augmentation settings by empirically evaluating properties of the transcriptome-wide predictions. The most relevant improvements were achieved by incrementally increasing the number of hard negatives in both the training set and each mini-batch. Increments were applied until no further motif or structural refinements were observed. The optimal settings were as follows:

- Sampled *N*_0_ hard negatives for Types I, III, and IV
- Sampled 2*N*_0_ hard negatives for Type II (NSUN6) to counteract the large number of false positives assigned to Type II m^5^C
- Hard negatives are selected uniformly across probability bins with an upper bound of 1.0
- Rebuilt train/validation splits so that the total negatives equaled the original positives plus *N*_0_ hard negatives for each of Types I/III/IV and 2*N*_0_ for Type II, giving an overall 6: 1 negative:positive ratio.
- Adjusted mini-batch sampling to 83% negatives and 17% positives (*∼*5:1 negative:positive ratio)
- Trained models from scratch (no pretrained weights) with early stopping on the rebalanced validation set

### 4.5 Transcriptome-wide predictions and functional enrichment

#### 4.5.1 Transcriptome-wide inference

All transcripts of GENCODE v45 (GRCh38) were scanned cytosine-by-cytosine. For each cytosine we generated a 51-nt window (*±* 25 nt). Windows that had been used during model training or testing were masked and excluded from downstream analyses.

#### 4.5.2 Top calls and gene sets

Writer-specific models produced probability scores for every cytosine. For each NSUN class we kept the 1 000 highest-scoring windows on Ensembl Canonical transcripts and collapsed the list to unique gene identifiers.

#### 4.5.3 Enrichment analysis

Gene lists were submitted to `g:Profiler`^63^ querying six resources: GO (BP, MF, CC), KEGG, Reactome and HPO. Benjamini–Hochberg FDR was applied; terms annotating *<* 10 or > 500 genes were discarded to avoid the instability of very small sets and the vagueness of very large ones. Finally, to remove largely overlapping annotations we applied a greedy Jaccard filter. Details on redundancy pruning and underlying rationale are provided in Suppl. Note 6.

### 4.6 Surrogate models for the interpretability of the oracle predictions

To interpret what our final Bi-GRU had learned, we trained surrogate models for each NSUN class using the MAVE–NN framework,^55^ which fits genotype–phenotype maps from multiplex assays of variant effects (MAVEs). Following Seitz et al.^54^ we generated an *in silico* MAVE dataset by treating the Bi-GRU as an oracle. Specifically, we: (i) collected transcriptome-wide windows predicted as methylated for each NSUN class; (ii) introduced random mutations at a 10% Poisson rate to generate *∼*1M variants per class; and (iii) rescored wild-type and mutated windows with the oracle to obtain predicted methylation probabilities. For each class, we uniformly sampled 100,000 sequences across probability bins to balance the training distribution.

Surrogate models were trained independently for the four NSUN classes using pairwise genotype–phenotype maps with a global epistasis nonlinearity and skewed-*t* noise model. We split the four datasets randomly into 90% for training and 10% for testing. Then, we further divided the training set into 90% for training and 10% for early stopping validation. Training employed Adam with learning rate 10^−4^, batch size 64, early stopping (patience 10), and default *L*_2_ regularization. See Suppl Notes 8-9 for further details.

## 5 Data and code availability

All resources are available at https://github.com/AnacletoLAB/RNA_m5C_predict, including:

- Training and evaluation datasets
- Python implementation of Bi-GRU, CNN and Transformer models
- Complete training and analysis code
- Pretrained model weights
- Transcriptome-wide predictions for GENCODE v45 (4 MB compressed)

Transcriptome-wide predictions are also archived at Zenodo https://doi.org/10.5281/zenodo.16629378in.xlsxformat.

Furthermore, to facilitate community use, we provide a user-friendly prediction tool at https://github.com/AnacletoLAB/RNA_m5C_predict. This standalone Python script accepts FASTA-formatted RNA or DNA sequences, automatically handles sequence preprocessing (including U-to-T conversion), and outputs writer-specific methylation probabilities for each cytosine. The implementation uses the final Bi-GRU model trained with heavy hard-negative augmentation and supports both CPU and GPU inference.

The software requires only Python, PyTorch, and Pandas, ensuring broad compatibility. For user convenience, the tool automatically handles ambiguous bases and sequence padding at transcript termini.

## 7 Author contributions statement

E.S. designed this research work, developed and implemented the AI and computational methods, conceived and executed the experiments, analyzed the results and contributed to the draft version of the paper. E.C., A.P. and G.V. contributed to the analysis of the results, to the overall structure of this work and drafted the paper. G.V. and E.C supervised the overall work. All the authors revised and approved the final manuscript.

## 8 Acknowledgments

This work was supported by National Center for Gene Therapy and Drugs Based on RNA Technology (MUR Project no. CN 00000041) funded by NextGeneration EU program, and by FAIR (Future Artificial Intelligence Research) project, funded by the NextGenerationEU program within the PNRR-PE-AI scheme (M4C2, Investment 1.3, Line on Artificial Intelligence) - AIDH – FAIR - PE0000013.

