## Supplementary Information for "AI methods and biologically informed data curation enable accurate RNA m^5^C prediction"

**Suppl. Note 6: Gene enrichment details** **10**

**Suppl. Note 7: Comparison with state-of-the-art m$^5$C predictors** **10**

**Suppl. Note 8: Model interpretability** **11**

**Suppl. Note 9: Surrogate model description** **12**

### Suppl. Note 1: m$^5$C detection limitations

In this section we introduce the different wet-laboratory techniques for m$^5$C detection, and summarize current limitations. These include bisulfite sequencing (BS-seq), m$^5$C-RNA immunoprecipitation (m$^5$C-RIP), methylation-dependent individual-nucleotide resolution crosslinking and immunoprecipitation (miCLIP), and 5-azacytidine–mediated RNA immunoprecipitation (Aza-IP).[1–4]

m$^5$C-RIP relies on antibody enrichment, thus it fails to achieve base-resolution m$^5$C detection and might suffer from potential non-specific antibody binding. miCLIP and Aza-IP can only identify m$^5$C sites targeted by specific methyltransferases; which requires their overexpression, limiting their applicability to cultured cells and potentially disturbing the endogenous m$^5$C methylome.

The most widespread and high-throughput technique is Bisulfite Sequencing (BS-seq) .BS-seq identifies m$^5$C sites indirectly, depending on the full conversion of cytosines (Cs) into uridines (Us). Hence, incomplete conversion (e.g., in structured regions) can result in false positive signals during m$^5$C detection.[5,6] Consequently, the detection threshold for BS-seq is generally established at either 10% or 20%,[5,7,8] which is comparable to the stoichiometry of mRNA m$^5$C sites, with a median of approximately 20%.[5,9,10] This leads to a substantial loss of m$^5$C signals during analysis, thus leading to limited RNA bisulfite maps of m$^5$C in mammals, with to adult tissues featuring only a few hundred confidently methylated sites. Additionally, the stringent reaction conditions in BS-seq cause extensive RNA degradation, making m$^5$C detection challenging in low-input samples and low-abundance RNAs. Furthermore, the complete conversion of cytosines (Cs) into uridines (Us) drastically reduces sequence complexity, introducing biases in library construction, lowering sequencing quality, and diminishing mappability, which impairs m$^5$C detection in RNAs with inherently low sequence complexity, such as chromatin-associated RNAs (caRNAs).

As a result, these intrinsic limitations significantly obstruct our comprehension of m$^5$C methylations, particularly in mRNAs, where the exact number of m$^5$C sites continues to be a subject of debate among studies.[5,6,8,9,11]

### Suppl. Note 2: Why training labels cover only four human m$^5$C writers

While the human genome encodes eight known m$^5$C methyltransferases, only four (NSUN1, NSUN2, NSUN5 and NSUN6) contribute to detectable signal in poly(A)+ RNA-seq datasets.

The excluded methyltransferases fall into three categories: (i) those confined to specific cellular compartments (e.g., mitochondria), (ii) those with extremely low activity on mRNAs, or (iii) those that exclusively target specific non-coding RNAs such as rRNAs or tRNAs. While these enzymes are functionally important in their respective context, they do not contribute to the regulatory m$^5$C patterns found in nuclear and cytoplasmic messenger RNAs that are the focus of this study.

Below we summarize the biological niche of every known human m$^5$C writer:

- DNMT2 (TRDMT1): Catalyzes C38 methylation in four cytosolic tRNAs. There is limited evidence of low mRNA methylation.[12]

- **NSUN1 (NOP2)**: Canonical target is 28S rRNA, but can methylate mRNA (Type IV) when nuclear-envelope integrity is compromised. Type IV sites emerge in nocodazole-treated HeLa cells, an experimental condition that mimics nuclear lamina breakdown during mitosis, and vanish upon NOP2 knock-down.[13]

- **NSUN2**: Principal mRNA writer; recognises a G-rich stem-loop (Type I) and installs thousands of marks across cell types. Knock-down eliminates ∼2,000 sites in HeLa cells and blocks ALYREF-mediated mRNA export.[9] During early embryogenesis NSUN2 decorates maternal transcripts en masse and is essential for timely cell-cycle progression.[10]

- NSUN3 and NSUN4: Confined to mitochondria, modify mt-tRNA$^{Met}$ (C34) and 12S/16S mt-rRNAs, respectively. While methylation has been detected on mitochondrial RNAs,[14] the small size of the mitochondrial transcriptome (37 genes) limits the availability of data for comprehensive motif discovery or model training. Moreover, because their activity is confined to organelle-encoded RNAs, these enzymes are largely irrelevant for studies of cytosolic or nuclear mRNA methylation.

- **NSUN5**: A nuclear/nucleolar MTases methyltransferase that can access mRNA already in the nucleus, but particularly in mitosis, when most nuclear content is released into the cytoplasm. iMVP clustering and over-expression experiments reveal a GUNGCCA motif (Type III) on transcripts from nocodazole-arrested HeLa cells and human oocytes.[13] The same motif has been detected by Guarnacci et al.,[15] that created a highly curated set of m$^5$C sites retrieved from several bisulfite sequencing experiments.

- **NSUN6**: Deposits Type II marks (CUC/UCCA loop); NSUN6 and NSUN2 account for the bulk of maternal-mRNA methylation across six animal phyla.[10,16] Furthermore, NSUN6 has been shown to be involved in the methylation of a cohort of mRNAs involved in cell migration,[17] promoting cancer. Importantly, loss-of-function mutations in

NSUN6 have been shown to cause an autosomal recessive neurodevelopmental disorder,[18] as well as have been linked to Alzheimer's Disease and brain injury,[19] hinting its importance in brain function.

- NSUN7: Selectively methylates enhancer RNAs (eRNAs)[20] and a limited number of mRNAs.[21] Because most eRNAs lack poly(A) tails and have short half-lives, they are underrepresented in poly(A)-selected libraries and often fall below the read-depth threshold required for reliable BS-seq detection. We examined the small set of enriched methylated transcripts (few dozen sites) reported by Ortiz et al.[21] in NSUN7-silenced cells, but were unable to identify a reproducible binding motif. This raises the possibility that at least some of the reported sites reflect false-positive BS-seq calls rather than *bona fide* NSUN7 substrates.

### Suppl. Note 3: Dataset preprocessing

#### Suppl. Note 3.1: Transcriptome mapping of human $m^5C$ sites

**Step 1: Discard unannotated and intronic calls.** The BS-seq protocol of Liu et al.[10] profiles mature RNA; cytosines lying outside annotated exons are therefore likely artifacts. After mapping sites to hg19 and overlapping with GENCODE v19, we removed (i) sites lacking any transcript annotation and (ii) intronic sites whose genomic coordinate does not intersect an exon of the retained isoform.

Furthermore, for each gene, we retained the isoform containing the greatest number of modifications (or, in case of a tie, the longest isoform), which is important when selecting unmodified cytosines.

**Step 2: Junction-distance filter for NSUN2.** NSUN2-mediated $m^5C$ deposition is linked to alternative splicing.[22] If a cytosine lies very close to a splice junction we cannot determine whether its flanks were intronic or exonic at the moment of methylation. We therefore retained NSUN2 (Type I) windows only if the modified C was $\geq$15 nt from the nearest exon–intron boundary of *any* isoform of that gene.

#### Suppl. Note 3.2: STREME algorithm and clustering

BS-sequencing procedures can lead to false-positive methylation calls.[8] A similar issue affects other RNA modification detection pipelines, such as $m^6A$-seq, where researchers often filter out sites not associated with the DRACH (D = not C; R = purine; H = not G) motif[1] during model training.[24,25] However, this approach can omit rare sites represented by other motifs.[26] A more sensitive and unbiased strategy is to use motif discovery tools like MEME[27] or STREME[28] to identify distinct motif clusters and filter out false positives. Notably, Liu et al.[13] reported that single-pass MEME[27] or STREME[28] runs could not recover all four motifs, unlike their bespoke pipeline. We attempted to replicate their method but encountered a sensitivity–specificity trade-off: we could either recover all four clusters with noisy boundaries and false positives inclusion, or reduce false positives at the expense of cluster completeness and clarity. We also tested other distance-based clustering algorithms akin to iMVP (computing sequence-distance matrices), but these failed because motifs do not fully overlap and require alignment that can include uninformative regions. By contrast, tuning STREME to scan only the 51 nt with central cytosine and using the "Central Distance" objective substantially improved motif detection.

Central Distance, during STREME enrichment test, utilizes the Bates distribution[29] to score positional enrichment. Let $d_i$ be the absolute offset (in nts) of the best-scoring site of sequence $i$ from the sequence midpoint and $L$ the (common) sequence length. STREME rescales this to $u_i = 2d_i/L$, which is $\mathcal{U}(0,1)$ under the null of uniformly placed sites. For every log-likelihood threshold $\tau$ it keeps the $n_\tau$ sequences whose site score exceeds $\tau$ and forms the sample mean $\bar{u}_\tau = \frac{1}{n_\tau}\sum_{i=1}^{n_\tau} u_i$. Because $\bar{u}_\tau$ then follows a Bates($n_\tau$) distribution, STREME computes the one-sided p-value $p_\tau$ and selects the threshold $\tau^* = \arg\min_\tau p_\tau$. Sequences scoring below $\tau^*$ are discarded; the survivors realign the PFM in the next refinement step. This left-tail test strongly down-weights the flanks and forces the algorithm to concentrate on the central position, a design originally introduced for ChIP-seq peak sequences,[28] but that translates well to our centered motifs. Nonetheless, STREME's stochastic initialization can sometimes yield slightly different (or even missing) motifs. To mitigate this, we reran STREME nine times (three random seeds $\times$ three parameter settings). Since motif lengths can vary, we set the maximum motif length to 8, 9, or 10 nt for each seed. Minimum length was kept at 5. Additionally, because STREME may locate motifs at any sequence position, we discarded any sequences in which the detected motif did not align with its expected central location.

After rebuilding PFMs from the filtered sequences and rescoring the entire dataset, we lost a fraction of sites, predominantly from the Type I cluster. This likely reflects the inherently weaker definition of the Type I motif. Given that most original sequences clustered as Type I in Liu et al.,[13] it is plausible that many false positives also accumulated there, weakening the motif's specificity.

[1]This motif is the main consensus for $m^6A$ sites in human.[23]

### Suppl. Note 4: Pooling and classification head

Transformer encoder and Bi-GRU final representations were pooled by exploring six different methods. The final pooled vector of the Bi-GRU and the Transformer and the final flattened vector for the 1D-CNN are fed to a classification head shared across architecture for fair model comparison.

#### Suppl. Note 4.1: Pooling

To feed the sequence to the fully connected layers for classification we must collapse the output hidden matrix $H \in \mathbb{R}^{L\times d}$ to a single summary vector $\mathbf{c} \in \mathbb{R}^d$. Below we formalize the six pooling schemes evaluated in this work.

1. **Average pooling**

$$\mathbf{c}_{\text{avg}} = \frac{1}{L}\sum_{i=1}^{L}\mathbf{h}_i.$$

2. **Max pooling**

$$\left[\mathbf{c}_{\max}\right]_j = \max_{1\le i\le L} h_{i,j}, \qquad j = 1,\dots,d.$$

3. **Central vector** This method consists in selecting only the hidden state of the central cytosine $\mathbf{c}_{\text{central}}$.

$$\mathbf{c}_{\text{central}} = \mathbf{h}_c.$$

   Since the central position is always occupied by cytosine, the input embedding at this position is constant across all sequences. In this sense, the representation of the central base resembles the role of a CLassify Token ( `[CLS]` ): it starts from the same initial vector and integrates contextual information from the surrounding nucleotides. However, unlike `[CLS]`, is not a trainable placeholder.

4. **Weighted Pooling** For each hidden state $i \in L$:

$$\begin{aligned} e_i &= \mathbf{w}^\top \mathbf{h}_i + b, \\ \alpha_i &= \frac{\exp(e_i)}{\sum_{j=1}^{L}\exp(e_j)}, \end{aligned}$$

   Then each $\alpha_i$ is used as weight to pool the hidden representation as:

$$\mathbf{c}_w = \sum_{j=1}^{L}\alpha_j\,\mathbf{h}_j,$$

   with trainable parameters $\mathbf{w}\in\mathbb{R}^d$ and $b\in\mathbb{R}$.

5. **Central Weighting**

$$\begin{aligned} \mathbf{s} &= \mathrm{GELU}\big(W^\top \mathbf{h}_c + \mathbf{b}\big) \quad \in \mathbb{R}^L, \\ s_c &= -\infty, \\ \alpha_t &= \frac{\exp(s_t)}{\sum_{j\neq c}\exp(s_j)}, \\ \mathbf{c}_{\text{cent-att}} &= \sum_{j=1}^{L}\alpha_j\,\mathbf{h}_j, \end{aligned}$$

   where $W\in\mathbb{R}^{d\times L}$ and $\mathbf{b}\in\mathbb{R}^T$ are trainable. GELU is a Gaussian Error Linear Unit.[30] The central hidden state $\mathbf{h}_c$ thus queries the entire sequence and re-weights $\mathbf{H}$ before aggregation, while "self-querying" at the center itself is explicitly suppressed by setting $s_c = -\infty$.

6. **Attention Pooling**

$$\begin{aligned} Q &= H\,W_Q, \qquad K = H\,W_K, \qquad V = H\,W_V, \\ A &= \mathrm{softmax}\big(QK^\top/\sqrt{d}\big), \\ Z &= A\,V, \\ \mathbf{c}_{\text{att}} &= \frac{1}{L}\mathbf{1}^\top Z, \end{aligned}$$

where $H \in \mathbb{R}^{L\times d}$ is the hidden matrix, $W_Q, W_K, W_V \in \mathbb{R}^{d\times d}$ are learned projections, the softmax is taken row-wise, and $\mathbf{1} \in \mathbb{R}^L$ is an all-ones column vector that averages the rows of $Z$.

Each pooled vector $\mathbf{c}_\star$ is forwarded to the two-layer MLP described in Section 4.2 for five-way classification.

##### Suppl. Note 4.2: Classification head

A single Feed Forward Neural Network is shared across experiments for model comparison. The context vector is processed for multiclass classification as follows: $\mathbf{c} \to \text{Linear}(\max\{128, d_{\text{in}}/2\}) \to \text{GELU} \to \text{Dropout } 0.1 \to \text{Linear}(5)$. During training, these logits are directly passed to the cross-entropy loss, which internally applies a log-softmax. At inference, we apply a softmax to obtain normalized class probabilities that sum to one.

#### Suppl. Note 5: Hyperparameter tuning

To identify the optimal hyperparameters for each model (pooling strategies included), we conducted a grid search across the different architectures. To streamline the process and focus on meaningful configurations, we initially tested several hyperparameters to exclude those that hindered model convergence. The following paragraphs outline the rationale behind the specific hyperparameters we chose to fix. Consequently, we report the grid search for each model architecture.

**Scheduler.** We controlled the learning rate with `torch.optim.lr_scheduler.ReduceLROnPlateau`. If the validation Weighted AUPRC failed to improve for $n$ consecutive epochs, the scheduler halved the current learning rate, never allowing it to fall below $1 \times 10^{-7}$. We prefer this simple, metric-driven scheme over more elaborate policies because it behaves consistently across a wide range of hyperparameter settings.

**Optimizer.** Across the three architectures, during pretesting, we obtained similar performances on RNNs and 1D-CNNs utilizing Adam[31] with weight decay regularization (AdamW)[32] and Root Mean Squared propagation (RMSprop), so we decided to deploy both optimizers. For the transformers, we utilized only AdamW, since it is the most widespread optimizer for transformers, and they are more computationally demanding.

**Batch Sampler.** Due to the imbalanced classes in our dataset, we decided to utilize a probabilistic sampler for the generation of the data in each batch. In particular, since the negative class is more important than a single positive class, each mini batch draws a negative sample with 50% probability, the remaining probability is split equally across the 4 positive classes. The sampling distribution exactly matches the class weights used in the weighted AUPRC metric, ensuring that the model is trained under the same importance scheme as it is evaluated.

#### Suppl. Note 5.1: Bi-GRU hyperparameters

We first explored both Gated Recurrent Unit (GRU)[33] or Long Short Term Memory (LSTM).[34] By testing some hyperparameters combinations, we consistently found GRU to be better than LSTM, so we decided to proceed only by considering GRU. Most importantly, as expected, Bi-GRU was better than unidirectional GRU, since the former captures patterns from both side of the sequence. Therefore, we decided to proceed with Bi-GRU. Below, we show the hyperparameters tested for the Bi-GRU. Hidden and embedding configurations were only tested when the former was 2 or 3 times the latter.

| **Global settings** | |
|---|---|
| Batch size | {16, 32} |
| Learning rate | {1e$^{-4}$, 1e$^{-5}$} |
| Weight decay | {1e$^{-5}$} |
| Optimizer | {RMSProp, AdamW} |
| Scheduler | {ReduceLROnPlateau (patience = 4)} |
| Loss | {Cross Entropy} |
| Sequence length | {51} |
| $k$-mer size | {1} |
| **Model hyperparameters** | |
| Embedding dimension | {64, 128} |
| Hidden dimension | {128, 256, 384} |
| # layers | {1, 2, 3} |
| RNN cell | {GRU} |
| Bidirectional | {True} |
| Dropout | {0.1} |
| Pooling type | {Max, Average, Central Vector, Weighted, Central Weighting, Attention} |

#### Suppl. Note 5.2: Stacked 1D-CNN hyperparameters

Below we indicate the hyperparameters tested for the stacked 1D-CNN model. Only combinations where the number of filters, kernels and pooling operations corresponded to the number of layers are tested, since each operation occur once per convolutional block.

| **Global settings** | |
|---|---|
| Batch size | {16, 32} |
| Learning rate | {1e$^{-4}$, 1e$^{-5}$} |
| Weight decay | {1e$^{-5}$} |
| Optimizer | {RMSProp, AdamW} |
| Scheduler | {ReduceLROnPlateau (patience = 6)} |
| Scheduler | {ReduceLROnPlateau} |
| Loss | {Cross Entropy} |
| Sequence length | {51} |
| *k*-mer size | {1,2,3} |
| Encoding type | {One Hot, ENAC} |
| **Model hyperparameters** | |
| # layers | {[2, 3]} |
| Filters | {[32, 64, 128], [32, 64]} |
| Kernel sizes | {[3,3,3], [5,5,5], [3,3], [5,5]} |
| pooling type | Max pooling |
| Pool sizes | {[1,2,2], [1,2]} |
| Dropout | {0.2} |

#### Suppl. Note 5.3: Transformer encoder hyperparameters

**Feed-Forward Layers.** The hidden dimension $h_{\text{ffn}}$ of the FFN was kept a multiple of the model dimension $d_{\text{model}}$ across all configurations. In particular, it was expanded with the following formula:

$$h_{\text{ffn}} = \frac{2}{3}\, 4 \times d_{\text{model}}$$

The transition factor expands the FFN dimension 4 times the model dimension, so that the FFN scales with model size; while the factor of $\frac{2}{3}$ keeps the parameter count and FLOPs of the FFN the same as when using more classical nonlinear activations functions such as GeLU (as proposed by Shazeer et al.[35]).

**Attention dimension** The dimension of query, key and value is set to:

$$d_Q = d_K = d_V = d_{\text{model}}/n_{\text{heads}}$$

Below, we report the hyperparameters tested for the Transformer encoder.

| **Global settings** | |
|---|---|
| Batch size | {16, 32} |
| Learning rate | {1e$^{-4}$, 1e$^{-5}$} |
| Weight decay | {1e$^{-5}$} |
| Optimizer | {AdamW} |
| Scheduler | {ReduceLROnPlateau (patience = 6)} |
| Loss | {Cross Entropy} |
| Sequence length | {51, 101} |
| k-mer size | {1} |
| **Model hyperparameters** | |
| Embedding dimension | {400, 600} |
| # blocks | {2, 4} |
| # heads | {20} |
| Pooling head | {Max, Average, Central Vector, Weighted, Central weighting, Attention} |

#### Suppl. Note 5.4: Post-training and testing

After five-fold cross-validation, the top models for each architecture were retrained on the whole training set for a number of epochs equal to the average epochs measured during cross-validation. Since we no longer had a validation set, the learning was regularized using `torch.optim.lr_scheduler.ReduceLROnPlateau` if the training weighted AUPRC failed to improve for $n$ epochs on the training set (with $n$ set as previously). For the Transformer, due to its higher complexity and higher learning rate, we utilized cosine annealing with warm restarts[36] to avoid overfitting[2]. We then evaluated the performance of each architecture on the test set.

[2] with restarts every 8 epochs and minimum learning rate set at 1e-07.

### Suppl. Note 6: Gene enrichment details

**Why keep only the Ensembl Canonical isoform?** Restricting the analysis to the single *Ensembl Canonical* transcript per gene mitigates annotation noise and avoids overweighting genes that possess many low-confidence isoforms.

**Redundancy pruning with the Jaccard filter.** For two gene sets $A$ and $B$ their overlap is quantified as

$$J(A, B) = \frac{|A \cap B|}{|A \cup B|}.$$

Significant terms are traversed in ascending FDR order; a candidate term $T$ is retained only if $J(T, S) < \tau$ for every term $S$ already accepted, with the threshold set to $\tau = 0.5$. Empirically, this cut-off removes near-duplicate annotations while preserving biologically distinct terms, crucial for a clear visualization of the top-10 enrichments shown in the main figures. Fully unfiltered enrichment tables are provided at Zenodo `https://doi.org/10.5281/zenodo.16629378`, for the scientific community.

### Suppl. Note 7: Comparison with state-of-the-art m$^5$C predictors

**Deepm5C** Hasan et al.[37] encoded each input sequence using four different schemes, feeding them into eight classifiers (both deep-learning and traditional), for a total of 32 base models. They then selected the top eight classifiers and stacked their predicted probabilities as input to a final 1D-CNN meta-learner (a technique known as ensemble stacking). Crucially, all sequences were drawn from the (then-new) m$^6$A Atlas,[38] which provides single-nucleotide–resolution RNA modification data for multiple marks, including m$^5$C. For m$^5$C, the Atlas aggregates 95 391 sites from 26 experiments. After removing redundant 41-nt windows with CD-HIT at a 90% identity threshold, Deepm5C's authors assembled a benchmark set of 58 159 positive sequences (an order of magnitude larger than any prior dataset). Negatives were sampled at random from the genome (matching the positive count) rather than from transcriptomic contexts.

**MLm5c** Kurata et al.[39] also used an ensemble stacking approach, but with traditional machine-learning classifiers as base learners: Support Vector Machines (SVM), eXtreme Gradient Boosting (XGBoost), LightGBM, and Random Forest (RF). They trained each classifier on 11 different sequence-based encodings, comprising:

- **Composition-based descriptors**, which reduce each RNA window to a vector of $k$-mer frequencies (normalized by window length), capturing global compositional biases (e.g. mono-, di-, or tri-nucleotide counts) but ignoring positional information.
- **Position-based descriptors**, which for each fixed-length window and each position encode the nucleotide or $k$-mer present (via one-hot profiles or sliding-window statistics), thereby preserving spatial context and enabling the learning of positional motifs relative to the center.
- **Language-model embeddings**, which treat each $k$-mer as a "word" in a large RNA corpus and learn dense vector representations (word2vec) that capture the statistical context in which each $k$-mer appears.

From the multiple trained classifiers, they selected the top 20 by validation performance, then stacked their predicted probabilities as input features to a final logistic regression meta-classifier for the ultimate m$^5$C vs. non-m$^5$C prediction.

The authors of MLm5c report using the same benchmark dataset as Hasan et al.[37] However, to extend sequences from 41 to 201 nt, they applied an undisclosed preprocessing step that reduced both training and test set sizes.

Remarkably, several individual base learners based on composition-based encodings outperform the Deepm5C ensemble. The single best model is LightGBM trained on trinucleotide frequency encodings (TNC), achieving 91.5% accuracy on the test set. Here, a $k$-mer encoding represents the frequency of each $k$-nucleotide subsequence in an RNA window; for TNC ($k = 3$),

$$f(s) \;=\; \frac{N(s)}{L}, \quad s \in \{\mathrm{AAA}, \mathrm{AAC}, \ldots, \mathrm{UUU}\},$$

where $N(s)$ is the count of $k$-mer $s$ in the window and $L$ is the window length.

**Our Models** We trained our top Bi-GRU, stacked 1D-CNN and Transformer encoder architectures on both the Deepm5C[37] and MLm5c[39] datasets. We trained the models on each dataset separately, even though both originate from the m$^6$A Atlas, because MLm5c (which benefits from longer contexts) applied a remapping step that altered and reduced the original Deepm5C dataset. We preserved each study's original training and test splits, then further partitioned the training sets into 80% for training and 20% for early-stopping validation. Models were trained with random batching (since positives and negatives are balanced) and evaluated on the respective held-out test sets.

**LGBM Training.** We trained LightGBM with trinucleotide-frequency (TNC) encodings on both the MLm5c dataset[39] and our own dataset. For MLm5c, sequences were 201 nt long, since the authors reported benefits from longer context (likely due to better capturing compositional differences between mRNAs and largely untranscribed or intronic RNAs). Accordingly, for our dataset we used 151 nt windows (the maximum length recovered during preprocessing). For MLm5c dataset, we retained the original training/test split, using 20% of the training data for early-stopping validation. For our dataset, we held out 15% for testing and used the remaining 85% for training, again reserving 20% of the training set for validation. As Kurata et al.[39] did, we implemented the classifier with the `lightgbm` (Python package `lightgbm` v3) library and the TNC encoding described above. Training used at most `num_boost_round`=1000 boosting iterations, with early stopping after 100 rounds without AUC improvement on the validation set. All other hyperparameters were left at their default values; the most important ones are listed here for reproducibility:

- `boosting_type`=`gbdt`, `objective`=`binary`, `metric`=`auc`;
- `learning_rate`=0.1, `num_leaves`=31, `max_depth`=−1 (unlimited);
- `min_data_in_leaf`=20, `min_sum_hessian_in_leaf`=0.001;
- `feature_fraction`=`bagging_fraction`=1.0 (subsampling disabled), `max_bin`=255;
- `verbosity`=−1 and `seed`=123.

The parameters are identical to those used by Kurata et al.[39] The trained LGBM on MLm5c dataset obtained an accuracy of 91.6%, confirming Kurata et al. results as well as our correct implementation of the model.

To train LightGBM on our negatives versus MLm5c negatives, we first randomly sampled negatives from our dataset to match the count in the MLm5c set. Importantly, we had to remove padded sequences since the full dataset of Kurata et al.[39] does not have a single padded sequence. We then trimmed all MLm5c sequences to 151 nt to align with our maximum window length. For both datasets, 15% of sequences were held out for testing; the remaining 85% were split 80%/20% for training and early-stopping validation, respectively. All LightGBM hyperparameters were kept as previously.

**$m^6A$ Atlas predictions.** To predict methylated sites across the $m^6A$ Atlas, we re-trained our top multiclass Transformer (selected via grid search) on windows trimmed to 41 nt to match the Atlas sequences. No further hard-negative fine-tuning was performed, ensuring fairness by preventing the model from becoming overly stringent in positive selection.

### Suppl. Note 8: Model interpretability

To decipher what our deep network had actually learned, we trained a simpler surrogate model capable of approximating the patterns captured by our intricate nonlinear model. Surrogate models have inherently interpretable mathematical forms that approximate more complex models. We adopted the MAVE–NN framework,[40] a neural-network-based Python package that learns genotype–phenotype (G-P) maps from multiplex assays of variant effects (MAVEs), such as deep mutational scans. MAVE–NN G-P maps are conceptually similar to LIME;[41] however, MAVE–NN can capture global nonlinearities and heteroscedastic noise, both critical for modeling DNNs trained on noisy data such as MAVE (model details in Suppl. Note 9).

Lacking experimental mutagenesis data for our methylated sequences, we followed Seitz et al.[42] and generated an *in silico* MAVE dataset using the trained Bi-GRU as an oracle. Specifically, we (i) collected all transcriptome-wide methylated sequences predicted for each positive class, (ii) introduced random mutations into these sequences, and (iii) fed both wild-type and mutated windows to the oracle, which returned their predicted methylation probabilities. These three steps yielded a complete *in silico* MAVE dataset, where the inputs are natural and mutated sequences, and the phenotypes are the oracle's predictions, suitable for training the surrogate model.

**Pipeline.** We selected the best Bi-GRU model after heavy hard-negative training as the oracle. We predicted every cytosine across the transcriptome and divided the positively predicted sequences by class. We further removed sequences with left or right padding since the surrogate model expects sequences of the same length. Consequently, we mutated the sequences uniformly to reach a total of $n$ mutated sequences. The number of mutations $m$ for each sequence was drawn from a Poisson distribution with $\mathbb{E}[m] = \lambda = r \cdot l$, where $r$ is the mutation rate and $l$ is the length of the sequence. In particular, we set $r$ to 0.1 for all four positive classes and generated $n = 1$ million mutated sequences per class. We deliberately generated and predicted such a high number of sequences to create training sets spanning all probability values. Once we combined the mutated sequences with the natural ones, we divided the data into probability bins of 0.05 from 0 to 1, and sampled uniformly across bins (until a bin was empty) to obtain a dataset of $100,000$ sequences per class. At this point, we had four datasets, one per positive class, with probabilities spread across all ranges fairly uniformly. We split each dataset randomly into 90% for training and 10% for testing. Then, we further divided the training set into 90% for training and 10% for early stopping validation.

The surrogate models were then trained with MAVE–NN using a single, fixed set of hyperparameters. Each model treated the full 51-nt RNA sequence as input and employed a pairwise genotype–phenotype (G–P) map. Latent phenotypes were passed through the default monotonic, 50-node `nonlinear` global-epistasis (GE) function, after which measurement noise was modeled with a `skewed-t distribution` likelihood whose log-scale was a quadratic ($m = 2$) polynomial of the GE output. Optimization was performed with the Adam algorithm (learning_rate $= 1 \times 10^{-4}$, batch_size = 64); training ran for at most 100 epochs with early stopping (patience = 10) and automatic restoration of the best weights. The model is trained with an $L_2$ regularization with $\lambda_\theta$ and $\lambda_\eta$ kept as default (respectively $10^{-3}$ and $10^{-1}$). For clarity, $\theta$ corresponds to the parameters of the G-P map and $\eta$ corresponds to the parameters of the GE nonlinearity and the noise model. The same configuration was used for all four *in silico* MAVE datasets. Finally, the parameters were corrected with "Uniform" Gauge. Below, we report the number of sequences predicted across the transcriptome for each class, the number of mutated sequences, and the final cardinality datasets to train the surrogate model.

| Class | Initial Predictions | Mutated Sequences | Final Cardinality | Selected | Training | Validation | Test |
|---|---|---|---|---|---|---|---|
| Type I | 77,154 | 1,000,000 | 1,077,154 | 100,000 | 81,000 | 9,000 | 10,000 |
| Type II | 162,774 | 1,000,000 | 1,162,774 | 100,000 | 81,000 | 9,000 | 10,000 |
| Type III | 16,267 | 1,000,000 | 1,016,267 | 100,000 | 81,000 | 9,000 | 10,000 |
| Type IV | 87,920 | 1,000,000 | 1,087,920 | 100,000 | 81,000 | 9,000 | 10,000 |

### Suppl. Note 9: Surrogate model description

A deterministic G-P map $f(x)$ maps each sequence $x$ to a latent phenotype $\phi$. Global epistasis (GE) regression leverages ideas previously developed in the evolution literature by mapping the latent phenotype nonlinearly to a prediction $\hat{y}$, that represents the most probable measurement value. A noise model is then used to describe the distribution of likely deviations from this prediction. MAVE–NN supports heteroscedastic noise models based on three different classes of probability distribution: Gaussian, Cauchy, and skewed-t.

**Linear Model.** MAVE–NN, like LIME, assumes that the latent phenotype is given by a linear function $\phi(x; \theta)$ that depends on a set of G–P map parameters $\theta$. MAVE–NN supports four types of G–P map models, in our analysis we utilize the pairwise model. The pairwise model contains all parameters of the additive model:

$$\phi_{\text{additive}}(x; \theta) = \theta_0 + \sum_{l=0}^{L-1} \sum_{c} \theta_{l:c}\, x_{l:c}\,.$$

Here, given a sequence $x$, each position $x_{l:c}$ contributes independently to the latent phenotype through its additive coefficient $\theta_{l:c}$, which represents the increment (or decrement) in $\phi$ produced by placing base $c$ at position $l$. The pairwise model is instead given by

$$\phi_{\text{pairwise}}(x; \theta) = \phi_{\text{additive}}(x; \theta) + \sum_{l=0}^{L-2} \sum_{l'=l+1}^{L-1} \sum_{c,c'} \theta_{l:c,l':c'}\, x_{l:c}\, x_{l':c'}\,,$$

and includes also interactions between all pairs of positions. Note the convention of MAVE–NN requires $l' > l$ in the pairwise parameters $\theta_{l:c,l':c'}$.

**Nonlinearities.** GE (Global Epistasis) models assume that each measurement $y$ is a nonlinear function $g(\cdot)$ of the latent phenotype $\phi$, plus some noise. In MAVE–NN, this nonlinearity is represented as a sum of hyperbolic tangent sigmoids:

$$g(\phi; \alpha) = a \;+\; \sum_{k=0}^{K-1} b_k \, \tanh\bigl(c_k\, \phi + d_k\bigr)\,.$$

Here, $K$ specifies the number of hidden nodes contributing to the sum, and $\alpha = \{a, b_k, c_k, d_k\}$ are trainable parameters. The $K$ value utilized by MAVE–NN and in this work is 50. By default, MAVE–NN constrains $g(\phi; \alpha)$ to be monotonic in $\phi$ by requiring all $b_k \geq 0$ and $c_k \geq 0$.

**Noise model and training objective.** After the linear G–P map and global-epistasis (GE) nonlinearity, each sequence $x$ is assigned a deterministic mean prediction $\hat{y} = g\bigl(\phi(x; \theta)\bigr)$. The measurement $y$ is assumed to follow a noise model due to the inherent noise in MAVE datasets. We select Skewed Student–$t$ noise, with a quadratic polynomial estimation of the $\sigma$ parameter as a function of $\hat{y}$.

**Training loss.** The surrogate is trained by minimising the batch-averaged loss $\mathcal{L} = (1/N)\sum_{i=1}^{N} \ell_i$ with ADAM; gradients are obtained by standard back-propagation through the additive pairwise map, the GE tanh stack, and the scale function $\sigma(\hat{y})$. Given some theoretical assumption, this corresponds to maximising explainability gain of the oracle by the surrogate model. For further mathematical and theoretical details please refer to the work of Tareen et al.[40]

| | Number of Epochs | Weighted AUPRC | Macro AUPRC | Macro Precision | Macro Recall | Macro F1 | Macro AUROC | Accuracy |
|---|---|---|---|---|---|---|---|---|
| 1 | 43.2±4.4 | 0.974±0.001 | 0.968±0.002 | 0.892±0.006 | 0.953±0.007 | 0.920±0.006 | 0.992±0.001 | 0.936±0.005 |
| 2 | 45.8±8.5 | 0.973±0.002 | 0.967±0.004 | 0.894±0.007 | 0.952±0.005 | 0.920±0.006 | 0.992±0.001 | 0.938±0.003 |
| 3 | 41.6±18.8 | 0.973±0.002 | 0.967±0.002 | 0.895±0.006 | 0.948±0.010 | 0.919±0.004 | 0.992±0.001 | 0.938±0.002 |
| 4 | 31.6±10.4 | 0.972±0.002 | 0.966±0.003 | 0.902±0.006 | 0.944±0.006 | 0.921±0.004 | 0.992±0.001 | 0.935±0.006 |
| 5 | 35.4±14.4 | 0.972±0.003 | 0.966±0.004 | 0.893±0.009 | 0.951±0.008 | 0.920±0.007 | 0.992±0.001 | 0.937±0.003 |
| 6 | 25.4±5.0 | 0.972±0.002 | 0.966±0.003 | 0.899±0.006 | 0.946±0.009 | 0.920±0.006 | 0.992±0.001 | 0.937±0.004 |
| 7 | 36.4±11.4 | 0.972±0.003 | 0.966±0.005 | 0.890±0.012 | 0.953±0.006 | 0.918±0.006 | 0.992±0.001 | 0.937±0.004 |
| 8 | 40.8±6.1 | 0.972±0.003 | 0.966±0.004 | 0.893±0.008 | 0.946±0.006 | 0.917±0.007 | 0.992±0.001 | 0.935±0.002 |
| 9 | 45.2±6.8 | 0.972±0.003 | 0.966±0.005 | 0.891±0.006 | 0.950±0.005 | 0.917±0.003 | 0.992±0.001 | 0.936±0.003 |
| 10 | 22.2±6.2 | 0.972±0.001 | 0.966±0.001 | 0.902±0.005 | 0.945±0.005 | 0.922±0.003 | 0.992±0.000 | 0.938±0.004 |

Table S1: Bidirectional GRU average five-fold cross-validation performance of the grid-search top 10 models, ranked by Weighted AUPRC

| | Model | Batch Size | Embedding Type | K-mer Size | Learning Rate | Optimizer | Sequence Length | Weight Decay | Dropout Rate | Embedding Dimension | Hidden Dimension | Number of Layers | Pooling Type |
|---|---|---|---|---|---|---|---|---|---|---|---|---|---|
| 1 | RNN | 16 | One-Hot | 1 | 1e-05 | AdamW | 51 | 1e-05 | 0.100 | 128 | 256 | 3 | Central Weighting |
| 2 | RNN | 16 | One-Hot | 1 | 1e-05 | RMSProp | 51 | 1e-05 | 0.100 | 128 | 256 | 3 | Central Weighting |
| 3 | RNN | 16 | One-Hot | 1 | 1e-05 | RMSProp | 51 | 1e-05 | 0.100 | 128 | 384 | 3 | Central Weighting |
| 4 | RNN | 32 | One-Hot | 1 | 1e-04 | RMSProp | 51 | 1e-05 | 0.100 | 64 | 128 | 1 | Central Weighting |
| 5 | RNN | 16 | One-Hot | 1 | 1e-05 | AdamW | 51 | 1e-05 | 0.100 | 128 | 384 | 3 | Central Weighting |
| 6 | RNN | 32 | One-Hot | 1 | 1e-04 | AdamW | 51 | 1e-05 | 0.100 | 64 | 128 | 1 | Central Weighting |
| 7 | RNN | 32 | One-Hot | 1 | 1e-05 | AdamW | 51 | 1e-05 | 0.100 | 128 | 384 | 3 | Central Weighting |
| 8 | RNN | 16 | One-Hot | 1 | 1e-04 | AdamW | 51 | 1e-05 | 0.100 | 64 | 128 | 1 | Attention |
| 9 | RNN | 32 | One-Hot | 1 | 1e-05 | AdamW | 51 | 1e-05 | 0.100 | 128 | 384 | 3 | Central Vector |
| 10 | RNN | 32 | One-Hot | 1 | 1e-04 | RMSProp | 51 | 1e-05 | 0.100 | 64 | 128 | 3 | Central Weighting |

Table S2: Bidirectional GRU parameters of the top 10 models explored through grid search.

| | Number of Epochs | Weighted AUPRC | Macro AUPRC | Macro Precision | Macro Recall | Macro F1 | Macro AUROC | Accuracy |
|---|---|---|---|---|---|---|---|---|
| 1 | 22.6±11.1 | 0.973±0.002 | 0.968±0.002 | 0.889±0.022 | 0.951±0.004 | 0.917±0.012 | 0.991±0.001 | 0.936±0.003 |
| 2 | 26.2±10.5 | 0.973±0.002 | 0.967±0.003 | 0.899±0.010 | 0.948±0.008 | 0.922±0.006 | 0.992±0.001 | 0.936±0.005 |
| 3 | 23.6±5.9 | 0.973±0.001 | 0.967±0.001 | 0.878±0.010 | 0.954±0.003 | 0.912±0.007 | 0.992±0.001 | 0.936±0.004 |
| 4 | 16.6±4.3 | 0.972±0.001 | 0.966±0.002 | 0.892±0.016 | 0.951±0.009 | 0.918±0.007 | 0.992±0.001 | 0.939±0.004 |
| 5 | 23.8±6.5 | 0.972±0.002 | 0.966±0.003 | 0.892±0.010 | 0.954±0.004 | 0.920±0.007 | 0.991±0.001 | 0.937±0.003 |
| 6 | 25.0±9.3 | 0.972±0.002 | 0.966±0.003 | 0.889±0.012 | 0.952±0.006 | 0.918±0.005 | 0.991±0.001 | 0.937±0.004 |
| 7 | 16.6±7.9 | 0.972±0.003 | 0.966±0.004 | 0.880±0.012 | 0.953±0.003 | 0.912±0.008 | 0.991±0.001 | 0.936±0.005 |
| 8 | 14.2±2.9 | 0.972±0.003 | 0.965±0.005 | 0.890±0.002 | 0.952±0.010 | 0.918±0.004 | 0.992±0.001 | 0.936±0.005 |
| 9 | 16.8±1.9 | 0.972±0.001 | 0.965±0.002 | 0.882±0.006 | 0.955±0.003 | 0.914±0.004 | 0.992±0.001 | 0.937±0.005 |
| 10 | 15.0±6.2 | 0.972±0.002 | 0.965±0.002 | 0.897±0.014 | 0.947±0.012 | 0.920±0.007 | 0.991±0.001 | 0.939±0.004 |

Table S3: Transformer encoder average five-fold cross-validation performance of the grid-search top 10 models, ranked by Weighted AUPRC

| | Model | Batch Size | Embedding Type | K-mer Size | Learning Rate | Optimizer | Sequence Length | Weight Decay | Embedding Dimension | Pooling Type | Number Of Blocks | Number Of Heads |
|---|---|---|---|---|---|---|---|---|---|---|---|---|
| 1 | Transformer | 32 | One-Hot | 1 | 1e-04 | AdamW | 51 | 1e-05 | 600 | Attention | 4 | 20 |
| 2 | Transformer | 16 | One-Hot | 1 | 1e-04 | AdamW | 51 | 1e-05 | 400 | Weighted | 4 | 20 |
| 3 | Transformer | 32 | One-Hot | 1 | 1e-05 | AdamW | 51 | 1e-05 | 400 | Central Vector | 2 | 20 |
| 4 | Transformer | 16 | One-Hot | 1 | 1e-05 | AdamW | 51 | 1e-05 | 400 | Central Vector | 4 | 20 |
| 5 | Transformer | 16 | One-Hot | 1 | 1e-04 | AdamW | 51 | 1e-05 | 400 | Attention | 4 | 20 |
| 6 | Transformer | 16 | One-Hot | 1 | 1e-04 | AdamW | 51 | 1e-05 | 600 | Central Vector | 2 | 20 |
| 7 | Transformer | 16 | One-Hot | 1 | 1e-04 | AdamW | 51 | 1e-05 | 600 | Central Vector | 4 | 20 |
| 8 | Transformer | 32 | One-Hot | 1 | 1e-05 | AdamW | 51 | 1e-05 | 600 | Central Vector | 4 | 20 |
| 9 | Transformer | 16 | One-Hot | 1 | 1e-05 | AdamW | 51 | 1e-05 | 400 | Central Vector | 2 | 20 |
| 10 | Transformer | 32 | One-Hot | 1 | 1e-04 | AdamW | 51 | 1e-05 | 600 | Average | 4 | 20 |

Table S4: Transformer encoder parameters of the top 10 models explored through grid search.

| | Number of Epochs | Weighted AUPRC | Macro AUPRC | Macro Precision | Macro Recall | Macro F1 | Macro AUROC | Accuracy |
|---|---|---|---|---|---|---|---|---|
| 1 | 49.8±11.9 | 0.971±0.002 | 0.963±0.004 | 0.853±0.018 | 0.957±0.004 | 0.896±0.011 | 0.992±0.000 | 0.933±0.004 |
| 2 | 10.2±3.1 | 0.970±0.002 | 0.963±0.003 | 0.874±0.024 | 0.948±0.008 | 0.906±0.017 | 0.991±0.000 | 0.933±0.006 |
| 3 | 44.0±14.7 | 0.970±0.002 | 0.962±0.003 | 0.854±0.019 | 0.953±0.006 | 0.895±0.010 | 0.992±0.001 | 0.933±0.006 |
| 4 | 56.8±10.5 | 0.970±0.002 | 0.963±0.004 | 0.839±0.016 | 0.957±0.004 | 0.887±0.010 | 0.991±0.000 | 0.930±0.004 |
| 5 | 38.6±10.0 | 0.970±0.002 | 0.963±0.004 | 0.845±0.009 | 0.958±0.004 | 0.892±0.005 | 0.991±0.000 | 0.932±0.004 |
| 6 | 9.0±2.4 | 0.970±0.002 | 0.962±0.003 | 0.871±0.015 | 0.948±0.009 | 0.904±0.009 | 0.991±0.000 | 0.933±0.003 |
| 7 | 10.6±3.0 | 0.970±0.001 | 0.962±0.003 | 0.867±0.009 | 0.948±0.005 | 0.902±0.005 | 0.991±0.001 | 0.931±0.005 |
| 8 | 12.2±1.8 | 0.970±0.001 | 0.962±0.003 | 0.857±0.012 | 0.951±0.007 | 0.897±0.007 | 0.991±0.000 | 0.931±0.005 |
| 9 | 55.6±6.7 | 0.970±0.002 | 0.962±0.004 | 0.842±0.019 | 0.956±0.005 | 0.888±0.012 | 0.992±0.000 | 0.932±0.005 |
| 10 | 49.4±23.2 | 0.970±0.003 | 0.962±0.004 | 0.842±0.018 | 0.955±0.003 | 0.888±0.012 | 0.991±0.001 | 0.931±0.005 |

Table S5: Stacked 1D-CNN average five-fold cross-validation performance of the grid-search top 10 models, ranked by Weighted AUPRC

| | Model | Batch Size | Embedding Type | K-mer Size | Learning Rate | Optimizer | Sequence Length | Weight Decay | Dropout Rate | Kernels | Filters | Max Pooling | Layers |
|---|---|---|---|---|---|---|---|---|---|---|---|---|---|
| 1 | 1DCNN | 16 | One-Hot | 1 | 1e-05 | RMSProp | 51 | 1e-05 | 0.200 | [5, 5] | [32, 64] | [1, 2] | 2 |
| 2 | 1DCNN | 16 | One-Hot | 1 | 1e-04 | RMSProp | 51 | 1e-05 | 0.200 | [5, 5] | [32, 64] | [1, 2] | 2 |
| 3 | 1DCNN | 16 | One-Hot | 1 | 1e-05 | RMSProp | 51 | 1e-05 | 0.200 | [3, 3] | [32, 64] | [1, 2] | 2 |
| 4 | 1DCNN | 32 | One-Hot | 1 | 1e-05 | AdamW | 51 | 1e-05 | 0.200 | [5, 5] | [32, 64] | [1, 2] | 2 |
| 5 | 1DCNN | 16 | One-Hot | 1 | 1e-05 | AdamW | 51 | 1e-05 | 0.200 | [5, 5] | [32, 64] | [1, 2] | 2 |
| 6 | 1DCNN | 16 | One-Hot | 1 | 1e-04 | AdamW | 51 | 1e-05 | 0.200 | [5, 5] | [32, 64] | [1, 2] | 2 |
| 7 | 1DCNN | 32 | One-Hot | 1 | 1e-04 | RMSProp | 51 | 1e-05 | 0.200 | [5, 5] | [32, 64] | [1, 2] | 2 |
| 8 | 1DCNN | 32 | One-Hot | 1 | 1e-04 | AdamW | 51 | 1e-05 | 0.200 | [5, 5] | [32, 64] | [1, 2] | 2 |
| 9 | 1DCNN | 32 | One-Hot | 1 | 1e-05 | RMSProp | 51 | 1e-05 | 0.200 | [3, 3] | [32, 64] | [1, 2] | 2 |
| 10 | 1DCNN | 16 | One-Hot | 1 | 1e-05 | AdamW | 51 | 1e-05 | 0.200 | [3, 3] | [32, 64] | [1, 2] | 2 |

Table S6: Stacked 1D-CNN parameters of the top 10 models explored through grid search.

| | Sequence Length | Number of Epochs | Weighted AUPRC | Macro AUPRC | Macro Precision | Macro Recall | Macro F1 | Macro AUROC | Accuracy |
|---|---|---|---|---|---|---|---|---|---|
| 1 | 101 | 31.6±12.1 | 0.973±0.003 | 0.968±0.004 | 0.888±0.012 | 0.951±0.009 | 0.916±0.010 | 0.991±0.001 | 0.935±0.008 |
| 2 | 51 | 22.6±11.1 | 0.973±0.002 | 0.968±0.002 | 0.889±0.022 | 0.951±0.004 | 0.917±0.012 | 0.991±0.001 | 0.936±0.003 |
| 3 | 51 | 26.2±10.5 | 0.973±0.002 | 0.967±0.003 | 0.899±0.010 | 0.948±0.008 | 0.922±0.006 | 0.992±0.001 | 0.936±0.005 |
| 4 | 51 | 23.6±5.9 | 0.973±0.001 | 0.967±0.001 | 0.878±0.010 | 0.954±0.003 | 0.912±0.007 | 0.992±0.001 | 0.936±0.004 |
| 5 | 101 | 14.2±5.3 | 0.972±0.001 | 0.966±0.002 | 0.890±0.013 | 0.947±0.009 | 0.916±0.006 | 0.992±0.001 | 0.932±0.004 |
| 6 | 51 | 16.6±4.3 | 0.972±0.001 | 0.966±0.002 | 0.892±0.016 | 0.951±0.009 | 0.918±0.007 | 0.992±0.001 | 0.939±0.004 |
| 7 | 51 | 23.8±6.5 | 0.972±0.002 | 0.966±0.003 | 0.892±0.010 | 0.954±0.004 | 0.920±0.007 | 0.991±0.001 | 0.937±0.003 |
| 8 | 101 | 23.0±6.9 | 0.972±0.003 | 0.966±0.003 | 0.887±0.011 | 0.950±0.012 | 0.914±0.007 | 0.991±0.001 | 0.934±0.007 |
| 9 | 51 | 25.0±9.3 | 0.972±0.002 | 0.966±0.003 | 0.889±0.012 | 0.952±0.006 | 0.918±0.005 | 0.991±0.001 | 0.937±0.004 |
| 10 | 51 | 16.6±7.9 | 0.972±0.003 | 0.966±0.004 | 0.880±0.012 | 0.953±0.003 | 0.912±0.008 | 0.991±0.001 | 0.936±0.005 |

Table S7: Transformer average five-fold cross-validation performances by the top 10 models, considering both sequence lengths of 51 and 101 nt, of the grid search, ranked by Weighted AUPRC

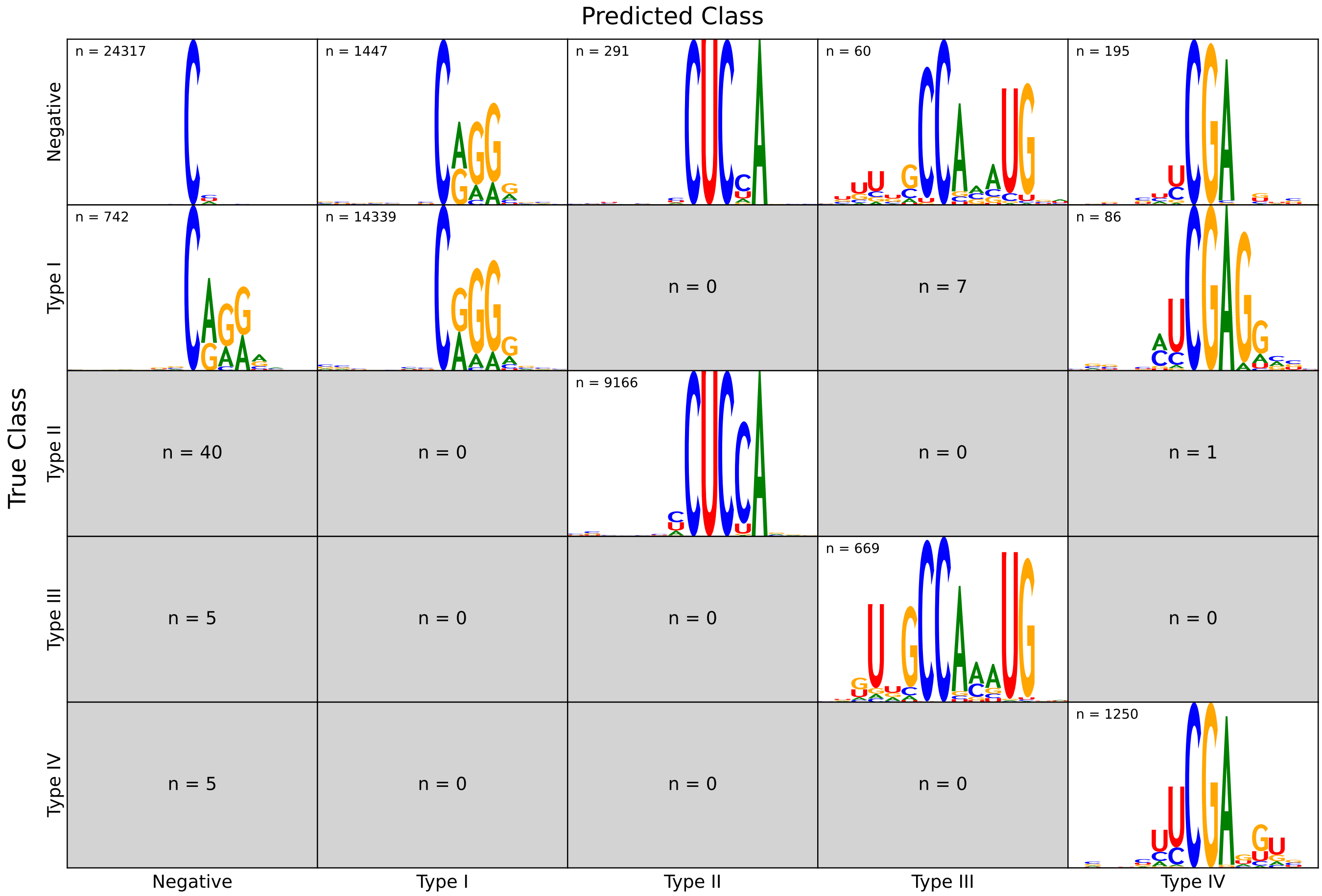

Figure S1: Performance of the final Transformer encoder on the combined training + test corpus. Rows are the true classes, columns the model's predictions. Inside each cell we report: (i) the number of windows assigned to that (true, predicted) pair; and (ii) a sequence-logo summarizing the corresponding windows whenever $n \geq 50$. Logos that would be based on fewer than 50 sequences are suppressed to avoid noisy motif representations.

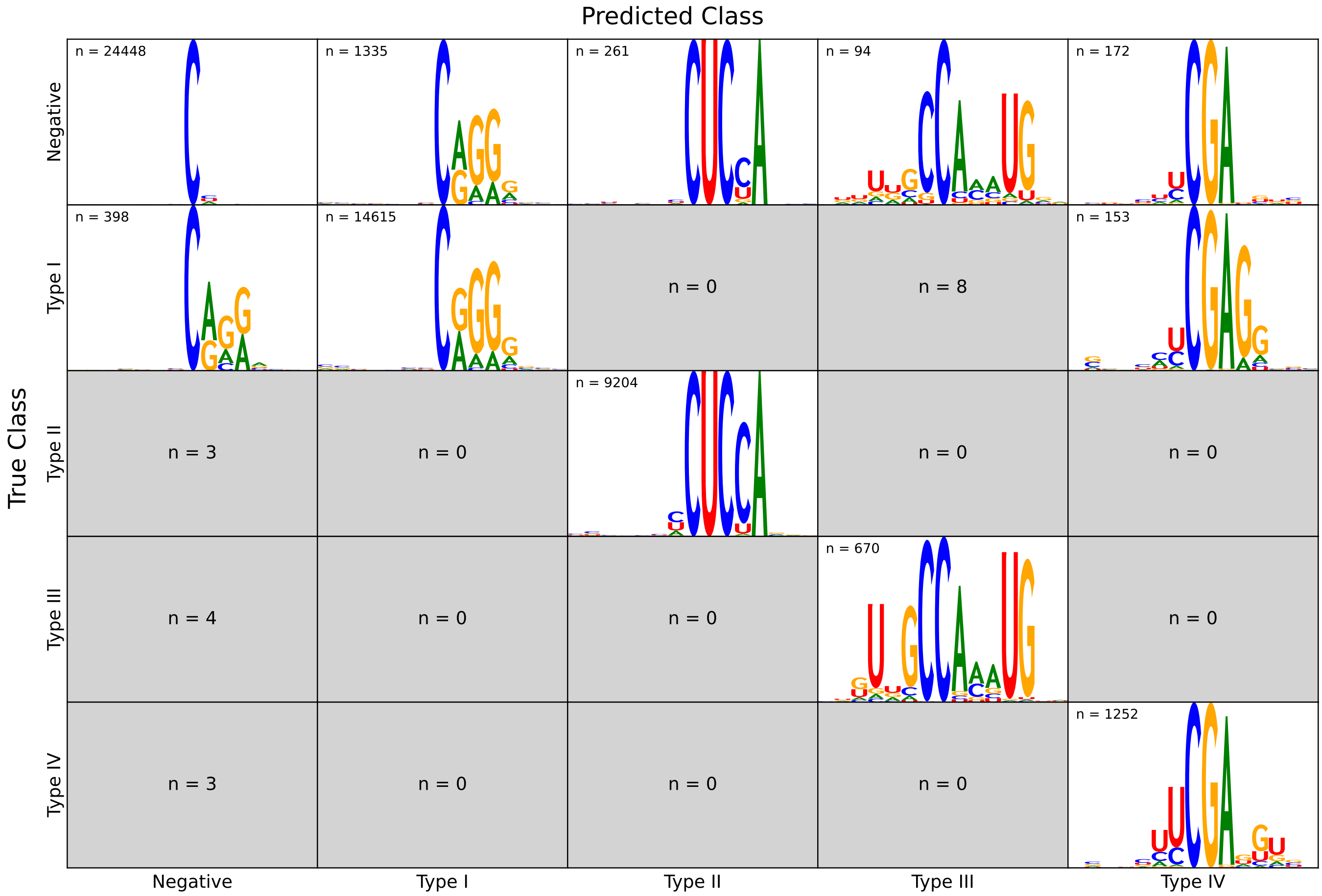

Figure S2: Performance of the final stacked 1D-CNN on the combined training + test corpus. Rows are the true classes, columns the model's predictions. Inside each cell we report: (i) the number of windows assigned to that (true, predicted) pair; and (ii) a sequence-logo summarizing the corresponding windows whenever $n \geq 50$. Logos that would be based on fewer than 50 sequences are suppressed to avoid noisy motif representations.

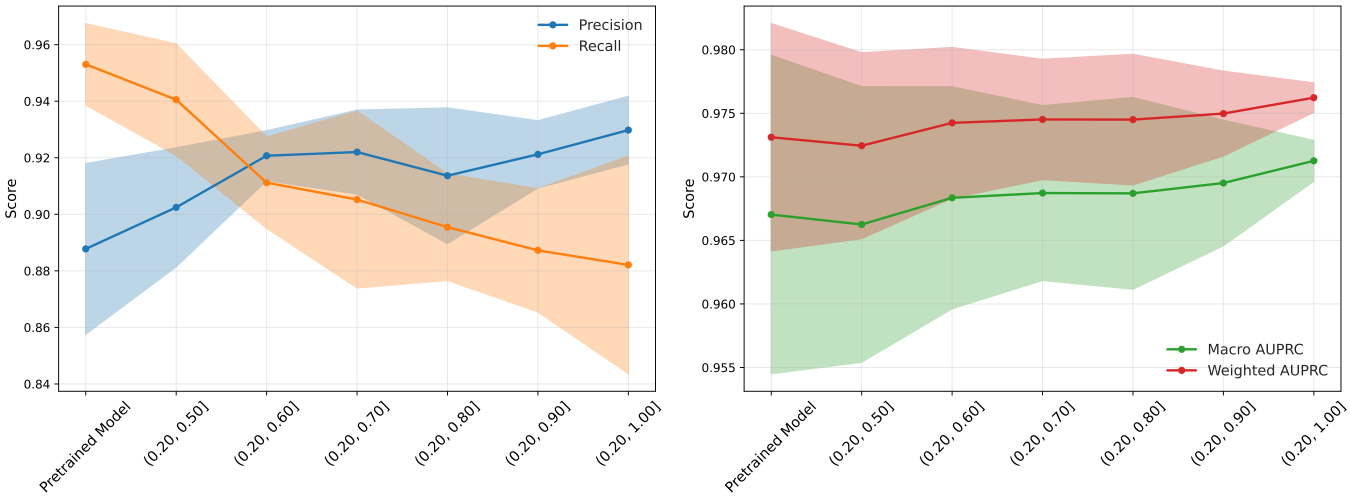

Figure S3: Average performance on the validation sets across folds for pretrained and lightly hard negative augmented models with different probability upper bounds for hard negative selection.

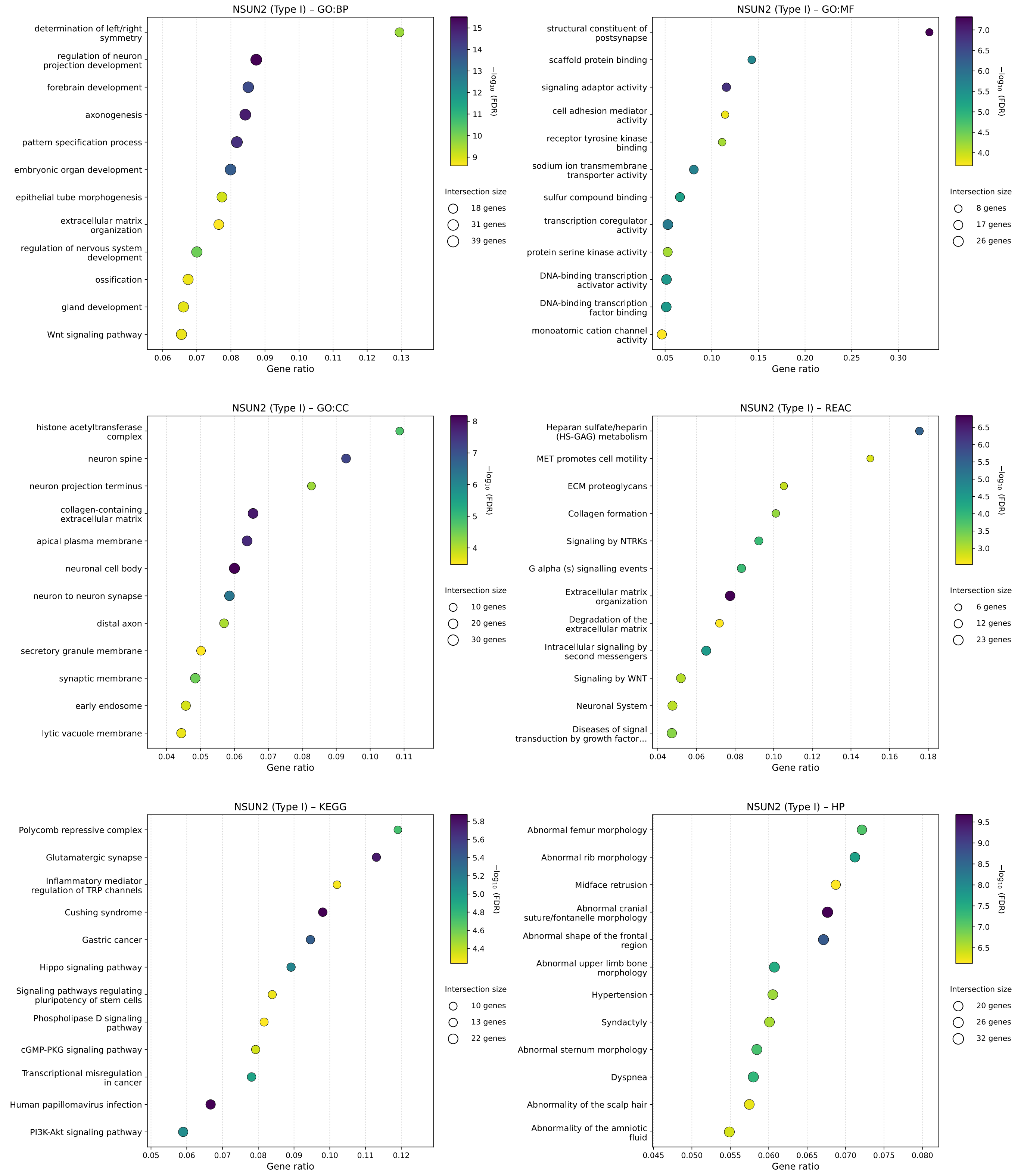

Figure S4: Enrichment analysis for transcripts predicted to be methylated by NSUN2 (Type I). Gene-databases are: Gene Ontology Biological Process (GO:BP), Gene Ontology Molecular Function (GO:MF), Gene Ontology Cellular Component (GO:CC), Reactome (REAC), Kyoto Encyclopedia of Genes and Genomes (KEGG), Human Phenotype Ontology (HP).

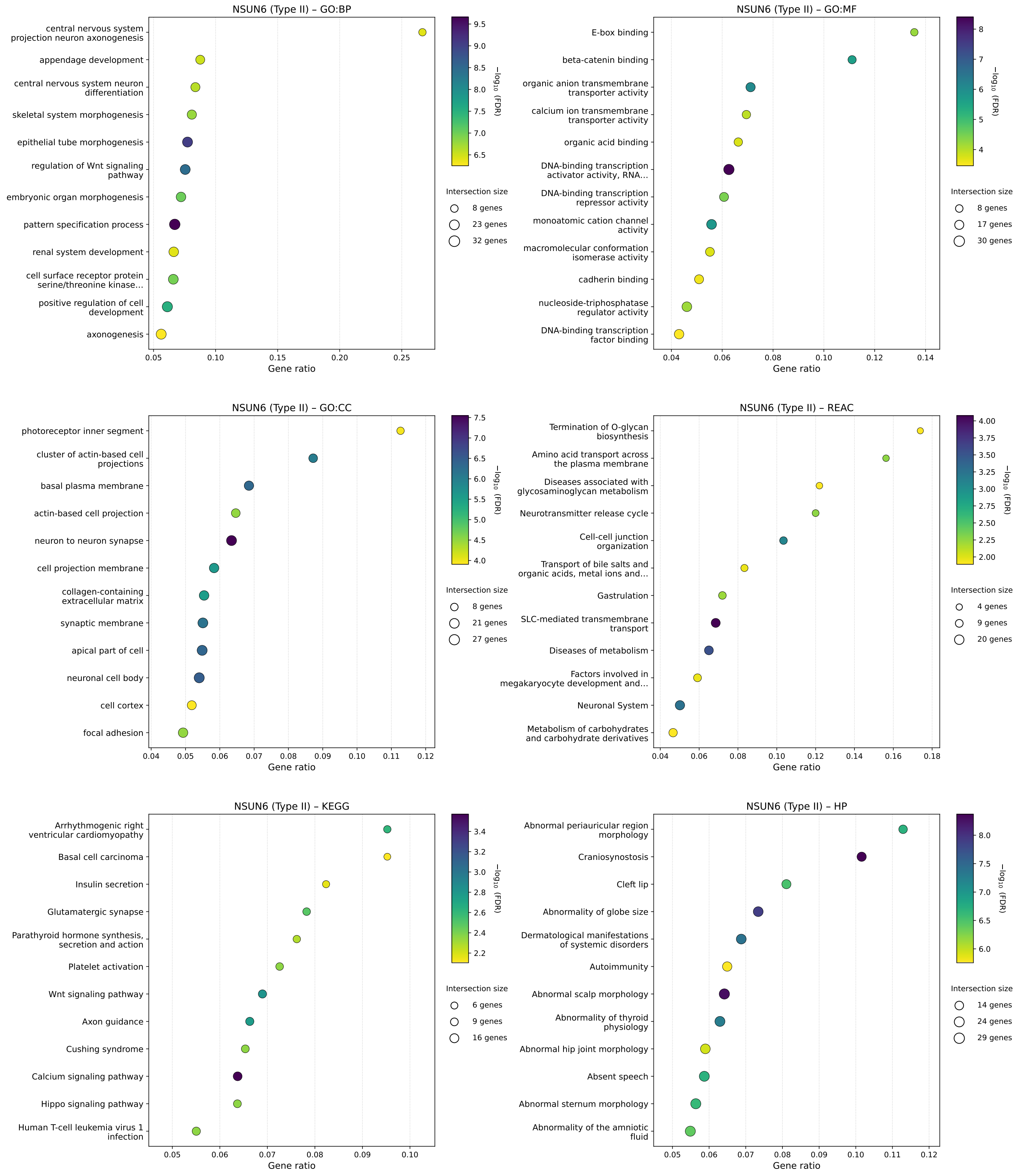

Figure S5: Enrichment analysis for transcripts predicted to be methylated by NSUN6 (Type II). Gene-databases are: Gene Ontology Biological Process (GO:BP), Gene Ontology Molecular Function (GO:MF), Gene Ontology Cellular Component (GO:CC), Reactome (REAC), Kyoto Encyclopedia of Genes and Genomes (KEGG), Human Phenotype Ontology (HP).

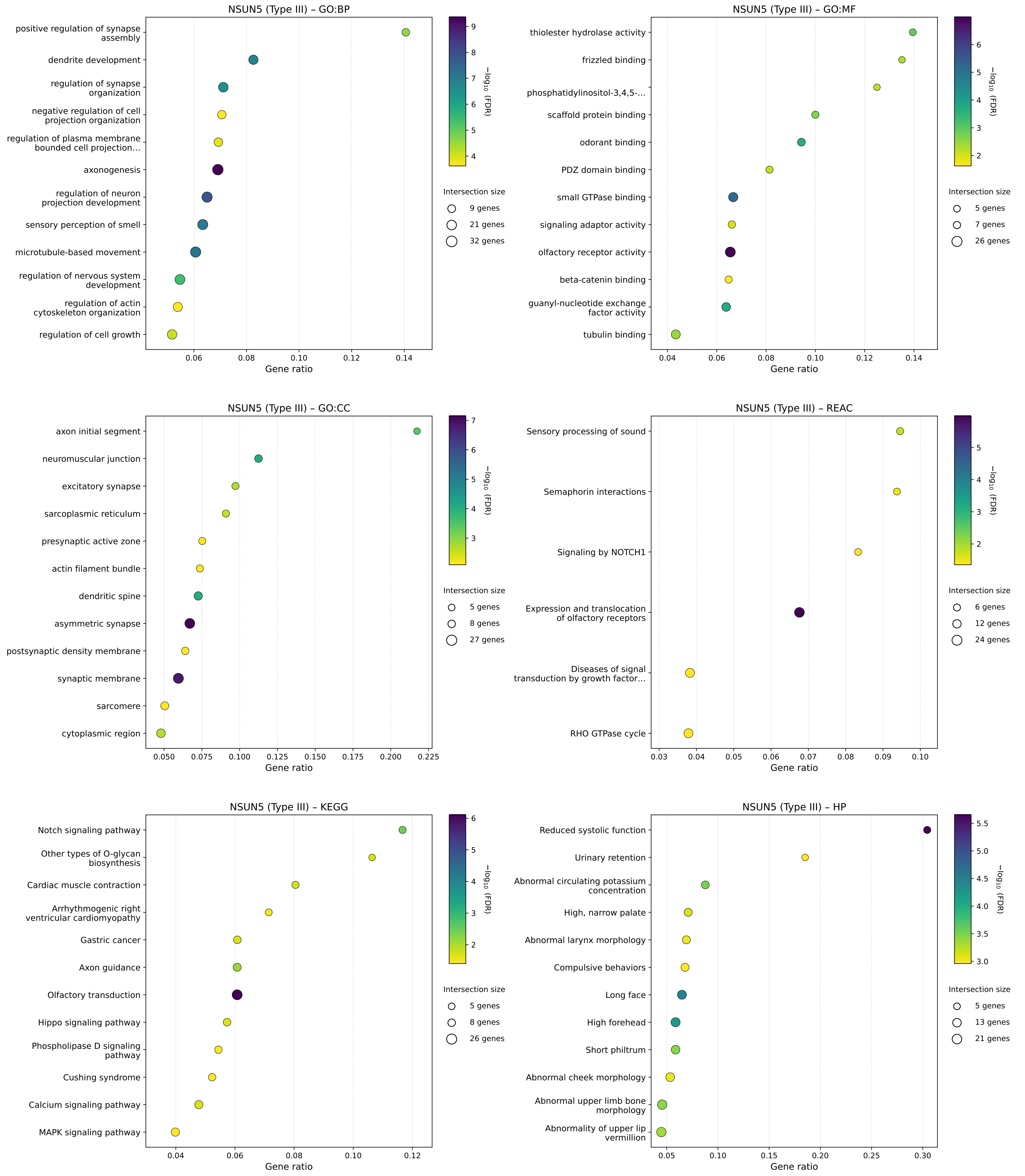

Figure S6: Enrichment analysis for transcripts predicted to be methylated by NSUN5 (Type III). Gene-databases are: Gene Ontology Biological Process (GO:BP), Gene Ontology Molecular Function (GO:MF), Gene Ontology Cellular Component (GO:CC), Reactome (REAC), Kyoto Encyclopedia of Genes and Genomes (KEGG), Human Phenotype Ontology (HP).

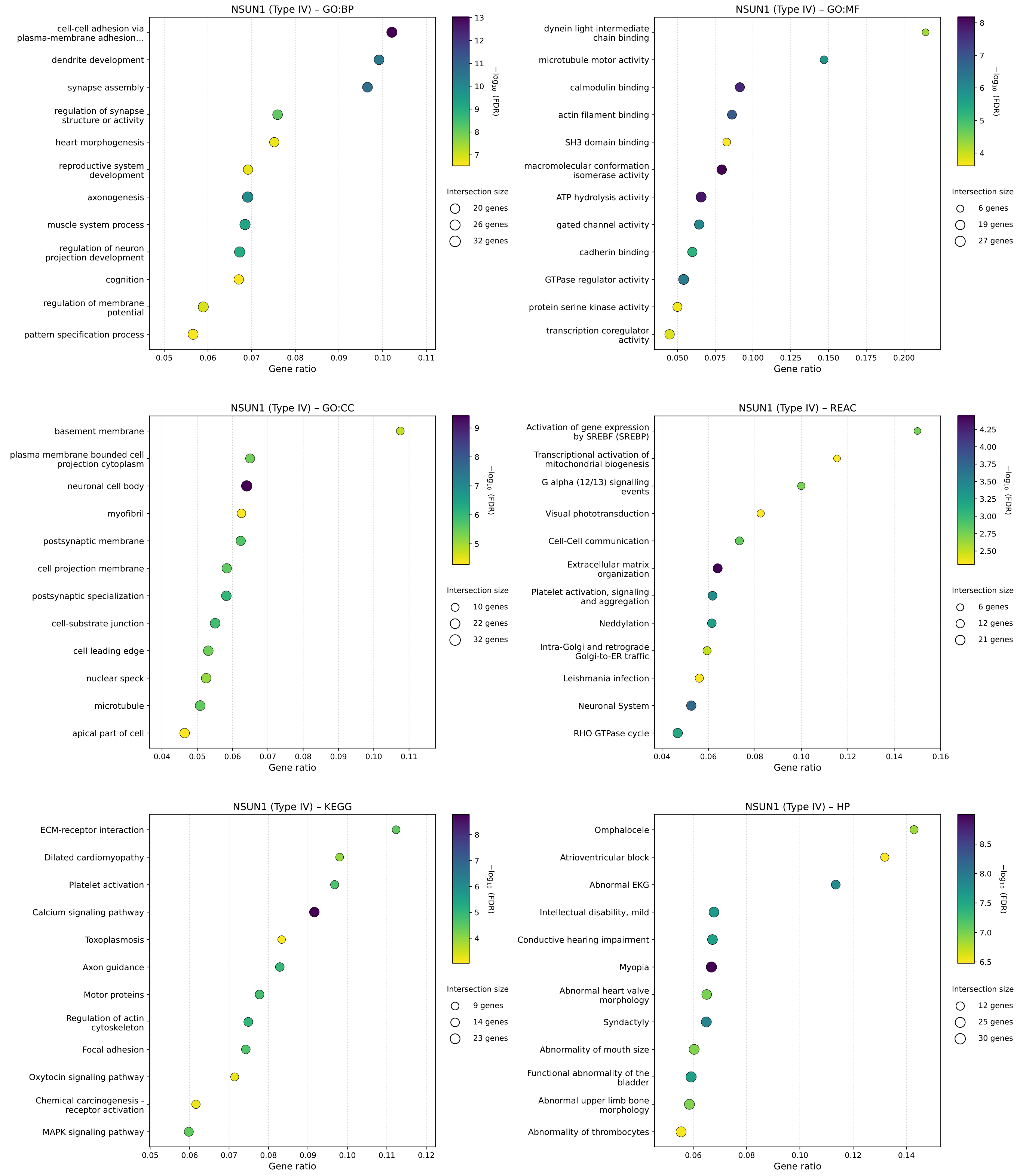

Figure S7: Enrichment analysis for transcripts predicted to be methylated by NSUN1 (Type IV). Gene-databases are: Gene Ontology Biological Process (GO:BP), Gene Ontology Molecular Function (GO:MF), Gene Ontology Cellular Component (GO:CC), Reactome (REAC), Kyoto Encyclopedia of Genes and Genomes (KEGG), Human Phenotype Ontology (HP).

Table S8: Performance comparison of Deepm5C, MLm5c and our models in binary classification.

| Dataset | Model | Accuracy | AUROC | AUPRC | Recall | Precision | F1 Score | Sensitivity | Specificity |
|---|---|---|---|---|---|---|---|---|---|
| Deepm5C | Bi-GRU | 0.848 | 0.916 | 0.927 | 0.819 | 0.869 | 0.844 | 0.819 | 0.877 |
| | 1D-CNN | 0.847 | 0.917 | 0.930 | 0.819 | 0.868 | 0.843 | 0.819 | 0.876 |
| | Transformer | 0.862 | 0.927 | 0.937 | 0.827 | 0.889 | 0.857 | 0.827 | 0.896 |
| | Deepm5C* | 0.852 | 0.938 | 0.901 | — | — | — | 0.846 | 0.857 |
| MLm5c | Bi-GRU | 0.870 | 0.937 | 0.940 | 0.850 | 0.885 | 0.867 | 0.850 | 0.890 |
| | 1D-CNN | 0.848 | 0.922 | 0.928 | 0.838 | 0.856 | 0.847 | 0.838 | 0.859 |
| | Transformer | 0.877 | 0.942 | 0.950 | 0.824 | 0.922 | 0.870 | 0.824 | 0.930 |
| | LGBM (TNC) | 0.916 | 0.971 | 0.976 | 0.880 | 0.949 | 0.913 | — | — |
| | MLm5c* | 0.939 | 0.977 | 0.978 | 0.941 | 0.937 | 0.939 | 0.941 | 0.938 |
| Current | Bi-GRU | 0.948 | 0.982 | 0.975 | 0.982 | 0.920 | 0.950 | 0.982 | 0.914 |
| | 1D-CNN | 0.931 | 0.976 | 0.970 | 0.973 | 0.897 | 0.934 | 0.973 | 0.888 |
| | Transformer | 0.946 | 0.981 | 0.974 | 0.984 | 0.915 | 0.948 | 0.984 | 0.909 |
| | LGBM (TNC) | 0.625 | 0.682 | 0.649 | 0.711 | 0.607 | 0.655 | — | — |

*Results taken from the authors' article

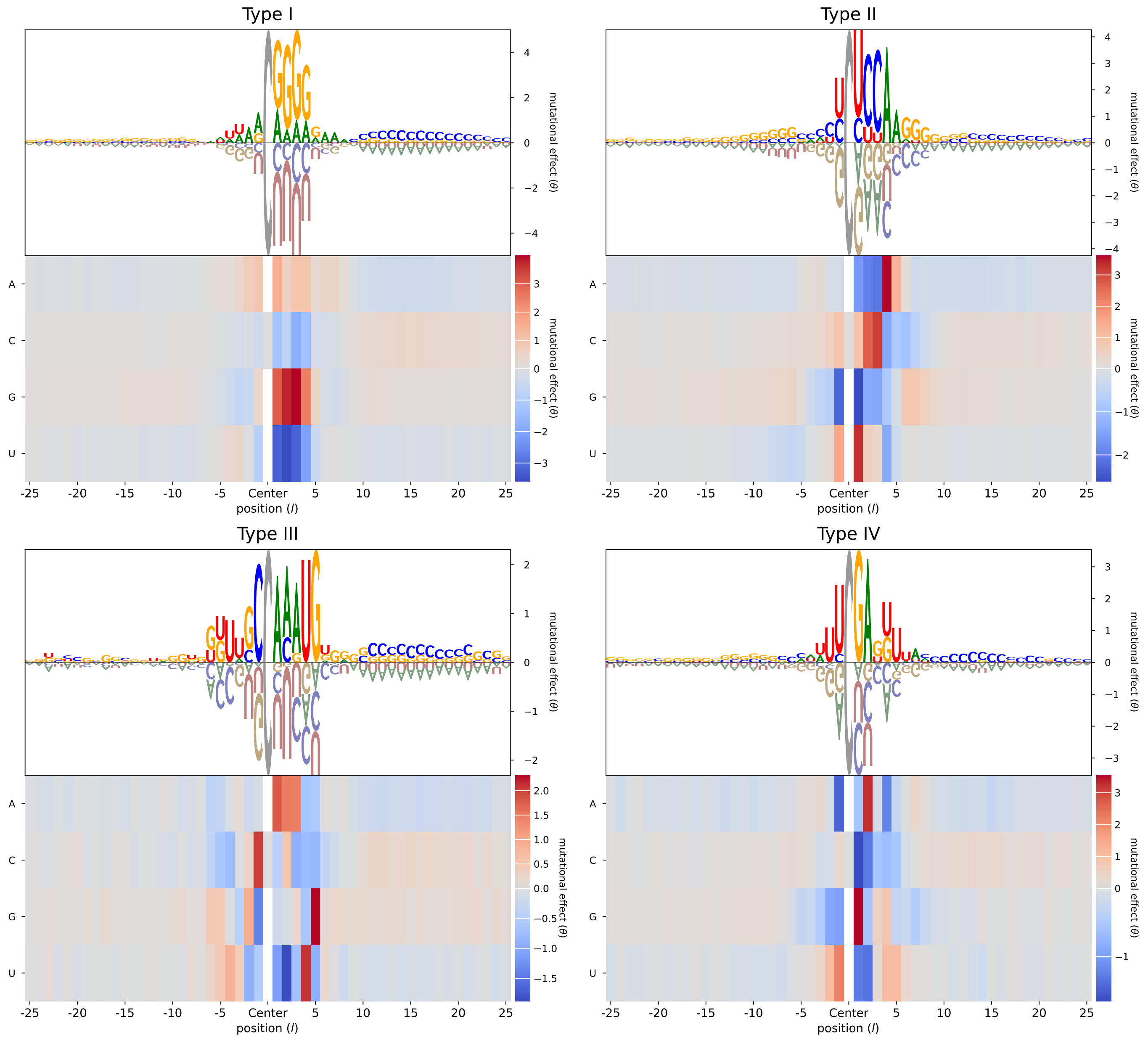

Figure S8: Additive effects captured by the surrogate models parameters $\theta_{l:c}$ trained with the mutational dataset constructed from each m$^5$C positive class. Larger positive $\theta_{l:c}$ raises the predicted modification probability, whereas large negative values lower it.

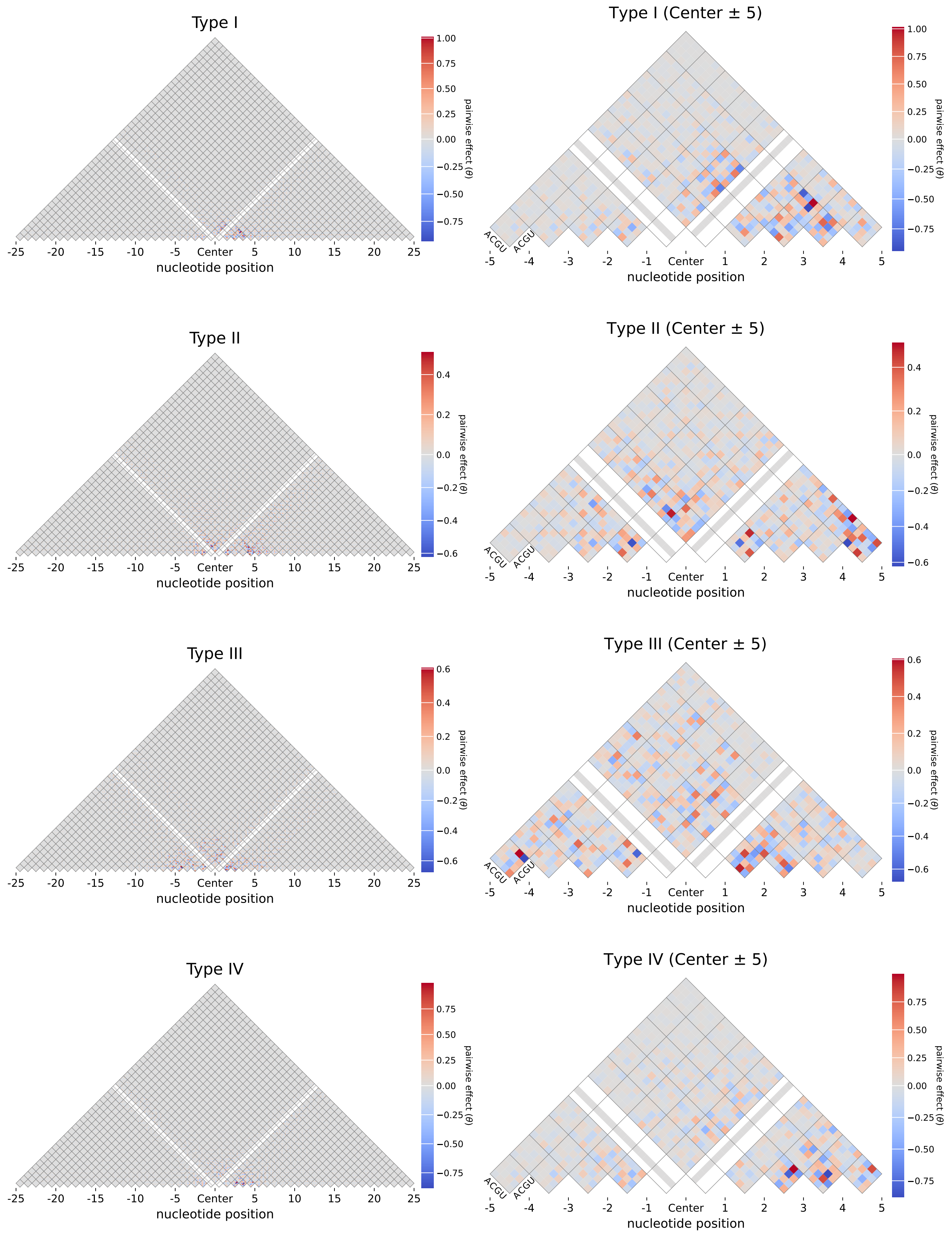

Figure S9: Pairwise (epistatic) effects captured by the surrogate models parameters $\theta_{l:c,\,l':c'}$ trained with the mutational dataset constructed from each m$^5$C positive class. Larger positive values raise the predicted co-occurring modification probability, whereas large negative values lower it.
